# Characterising the diffusion functional signature of negative BOLD with interleaved TMS-fMRI in the human brain

**DOI:** 10.64898/2026.03.20.713098

**Authors:** Inès de Riedmatten, Arthur P C Spencer, Roberto Martuzzi, Vincent Rochas, Jean-Baptiste Pérot, Filip Szczepankiewicz, Ileana O Jelescu

## Abstract

The coupling between brain excitatory activity and positive blood oxygen level-dependent (BOLD) responses is well-established. Although often associated with inhibition, negative BOLD remains partially understood. Moving away from neurovascular coupling, apparent diffusion coefficient (ADC)-fMRI provides a more direct measure of excitatory activity, possibly mediated by transient cellular deformations. While decreases in ADC align with positive BOLD, the possible translation of negative BOLD into positive ADC has not been investigated in humans. Diffusion-weighted fMRI (dfMRI) combines vascular and microstructural contributions. Using interleaved subthreshold transcranial magnetic stimulation (TMS)-fMRI on the primary motor cortex (M1), we induced negative BOLD responses in contralateral M1 and primary somatosensory cortex (S1). This was accompanied by a negative dfMRI response, but no ADC-fMRI response, indicating minimal microstructural fluctuations. In ipsilateral M1/S1, no BOLD response was detected while dfMRI revealed a positive cluster, suggesting sensitivity to subtle neural activity. These findings provide new insights into vascular and neuronal responses underlying subthreshold TMS and negative BOLD.

## 1 Introduction

Brain activity is commonly investigated using blood oxygen level-dependent (BOLD)-fMRI contrast, which reflects neuronal activity indirectly through neurovascular coupling [1]. In task-based fMRI, positive BOLD responses are generally interpreted as markers of neural excitation, defined as a transient increase in the probability of firing in a target cell [2, 3]. Conversely, negative BOLD responses may reflect neural inhibition, defined as a transient decrease in the probability of firing in a target cell [3–5]. Recently, apparent diffusion coefficient (ADC)-fMRI has emerged as a promising functional imaging alternative that links neuronal activity to morphological deformations through so-called neuromorphological coupling, offering improved spatial and temporal specificity relative to the neurovascular BOLD contrast [6, 7]. By leveraging sensitivity to water displacement, ADC-fMRI probes activity-related microstructural changes. While excitatory neuronal activity is typically associated with a decrease in ADC [6–10], likely driven by transient microstructural swelling, the ADC-fMRI signature corresponding to negative BOLD responses has yet to be characterised.

### 1.1 Blood oxygen level-dependent fMRI

BOLD-fMRI is sensitive to changes in the ratio of oxy-to deoxyhaemoglobin [11]. During excitatory brain activity, the marked increase in CBF exceeds the concomitant increase in cerebral metabolic rate of oxygen consumption (CMRO_2_). This results in a decrease in the oxygen extraction fraction (OEF) and in deoxyhaemoglobin concentration, captured as an increase in the measured 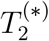-weighted MRI signal (positive BOLD) [2, 12, 13].

While positive BOLD is widely acknowledged to result from excitatory activity, the origins of negative BOLD remain unclear. There is substantial evidence that negative BOLD signals are associated with inhibitory neural activity, as shown with local field potential (LFP) and multi-unit activity (MUA) recordings [14, 15]. It has been reported that vasoconstriction was correlated to inhibitory synaptic activity, and in some cases, to a reduction in firing rate [15]. Vasoconstriction would translate into a decrease in CBF with a lesser reduction in CMRO_2_, which would in turn result in an increased OEF [5, 16]. Conversely, it has been suggested that negative BOLD responses may also arise from large increases in neuronal activity [17, 18], such as during seizures or during the initial dip of the BOLD response. In this case, CMRO_2_ exceeds the increase in CBF, resulting in insufficient oxygen supply. Although less likely, a decrease in CBF due to vascular “steal” by adjacent activated regions is often cited as a potential contributor to negative BOLD [19]. All of these mechanisms would result in a locally increased deoxyhaemoglobin concentration, a more heterogeneous magnetic field around vessels and thus a decrease in the measured 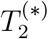-weighted MRI signal (negative BOLD). Despite the prevalent assumption that negative BOLD reflects inhibitory neuronal activity, the precise neurophysiological mechanisms underlying negative BOLD responses remain a matter of ongoing debate and active investigation [14].

### 1.2 Apparent diffusion coefficient fMRI

ADC timeseries are typically derived from pairs of diffusion-weighted images acquired with two distinct b-values, which, in combination with other acquisition design features, largely cancels out the *T*_2_-weighting and thus BOLD contributions to the ADC time courses [6, 7, 20]. The ADC timeseries are calculated according to Eq. 1,

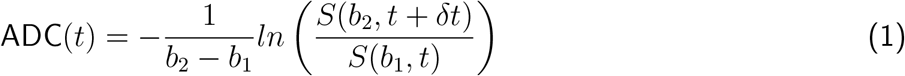

with 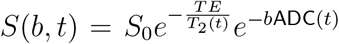. The two b-values are commonly acquired in an alternating fashion. The corresponding S(b_1_) and S(b_2_) timeseries can also be analysed individually and are referred to as diffusion fMRI (dfMRI). By combining T_2_- and diffusion-weighting, the dfMRI time courses retain partial BOLD-like contrast and sensitivity to microvasculature (via the spin-echo acquisition [21]) in addition to sensitivity to cellular microstructure (via diffusion-weighting). Diffusion-weighting also reduces direct contributions from intra-vascular signals.

While a functional decrease in ADC during excitatory brain activity has been linked to transient neuromorphological alterations, encompassing deformations and swelling in axons, cell bodies, dendrites, synaptic boutons and astrocytes processes, as shown in optical imaging studies [22–25], the neuromorphological changes associated with negative BOLD have not yet been investigated.

### 1.3 Transcranial magnetic stimulation

We used transcranial magnetic stimulation (TMS) combined with fMRI to modulate brain activity while simultaneously acquiring fMRI data, adopting a “perturb-and-measure” approach [26]. Previous studies have demonstrated that applying unilateral subthreshold repetitive TMS to the primary motor cortex (M1) at 3-5 Hz induces negative BOLD responses in the contralateral M1 [27–29], an effect that has also been implicated in supporting functional recovery in hemiparetic stroke patients [29]. In the context of motor function, this negative BOLD may reflect interhemispheric inhibition, which is thought to enable unilateral movement and high levels of manual dexterity in complex tasks [16].

The primary goal of this study is to investigate the ADC-fMRI response associated with negative BOLD, together with the responses of its constituent diffusion-weighted fMRI time courses, thereby providing new insights into the microstructural response underlying negative BOLD. As a secondary goal, because negative BOLD response is probably not simply a mirror of positive BOLD response but arises from distinct neurovascular coupling mechanisms [5], we also aim to use ADC-fMRI and dfMRI to disentangle neuronal contribution from vascular effects underlying negative BOLD signals.

## 2 Methods

### 2.1 Experiments

This study was approved by the ethics committee of the canton of Vaud, Switzerland (CER-VD). All participants provided written informed consent. Twelve healthy young adults (age 20-33 years, median 27; 2 females) were scanned. All subjects except one were right handed according to the Edinburgh Handedness Inventory [30].

Prior to the MRI scan, the right M1 hotspot was determined for each subject outside the scanner using single TMS pulses. The scalp location that consistently evoked muscle twitches in the left hand was marked with a cross. Once set up in the MRI scanner, the position of the MR-compatible TMS coil at the hotspot was confirmed and the rest motor threshold (RMT) was determined as the intensity eliciting identifiable motor-evoked potentials (MEPs) in half of the occurrences, as recorded from five pre-gelled electromyography (EMG) electrodes (EL508, BIOPAC Systems, Inc., Goleta, US) measuring the interosseus dorsalis primus manus and the abductor digiti minimi muscle groups of the left hand (Figure 1A).

**Figure 1.**
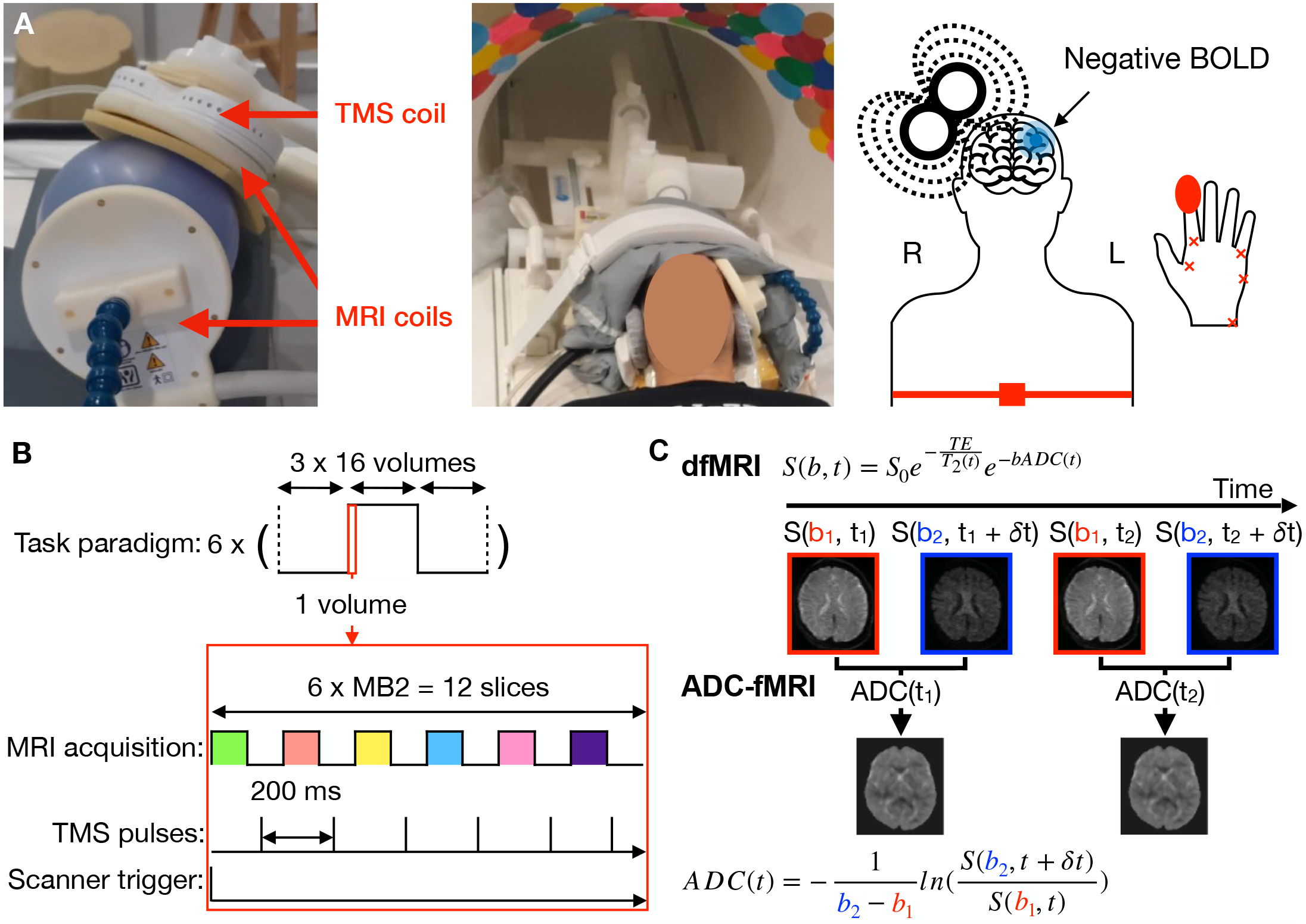
Experimental setup and MRI protocol. (A) TMS-fMRI setup. A combined TMS coil and seven-channel MRI coil were positioned over the participant’s right primary motor cortex. A second seven-channel coil was placed symmetrically over the left hemisphere. Surface electromyography (EMG) electrodes were placed on the left hand to localise the motor hotspot and determine the rest motor threshold. Other physiological signals were recorded using a respiration belt and a photoplethysmograph. (B) Interleaved TMS-fMRI acquisition used for both BOLD-fMRI and ADC-fMRI. The task consisted of six epochs of a baseline condition (19.2 s), followed by a TMS block (19.2 s) and a recovery period (19.2 s). During the TMS block, pulses were delivered at 5 Hz between the acquisition of MRI slices to prevent imaging artifacts. One image volume contained 12 slices, acquired with a multiband factor of 2. (C) Diffusion fMRI and ADC-fMRI acquisition. ADC time courses were derived by combining pairs of diffusion-weighted volumes acquired at two different b-values b_1_ (200 s mm^−2^) and b_2_ (1000 s mm^−2^). BOLD = blood oxygen level-dependent, ADC = apparent diffusion coefficient, dfMRI = diffusion functional MRI.

### 2.2 Physiological Monitoring

During the functional scans, photoplethysmography (TSD200-MRI), respiration belt (TSD201) (BIOPAC Systems, Inc., Goleta, US) and EMG data were recorded via a BIOPAC MP160 physiological monitoring system (BIOPAC Systems Inc, Goleta, California, USA). Triggers sent by the MRI scanner at every repetition time (TR) and triggers sent by the transcranial magnetic stimulator at every TR during the TMS blocks were also recorded via the BIOPAC system. AcqKnowledge data acquisition software (BIOPAC Systems Inc) was used to record signals sampled at 10 kHz. The cardiac and respiratory recordings were used to correct for physiological artifacts.

### 2.3 MRI Acquisition

MRI data were acquired using a 3T Siemens Magnetom Prisma with 80 mT/m gradients and 200 T/m/s slew rate. For anatomical reference and tissue segmentation, high-resolution T_1_-weighted images were acquired with a 64-channel head coil, using 3D Magnetization Prepared Rapid Acquisition Gradient Echoes (MPRAGE) with the following parameters: 1 mm^3^ isotropic voxels; 192 × 240 × 256 mm^3^ field of view (FOV); 256 slices; TR 2300 ms; echo time (TE) 2.98 ms; inversion times (TI) 900 ms; flip angle 9°.

A MR-compatible MRi-B91 Air Cooled TMS coil (MagVenture A/S, Farum, Denmark) was positioned tangential to the scalp of the subjects on the right M1 for TMS. A 7-channel (7ch) surface coil compatible with TMS (ALSIX GmbH, Vienna, Austria [31]) was positioned between the TMS coil and the scalp to receive the MRI signal (Figure 1A). Another 7ch surface coil was positioned symmetrically on the left M1.

For intermediate registration of the fMRI volumes to high-resolution anatomical T1-weighted images, two anatomical balanced steady state free precession (bSSFP) volumes were acquired: one with the body coil (BC); one with the 7ch coils, with the following parameters: (2 mm)^3^ isotropic voxels; 192 × 256 × 256 mm^3^ FOV; 128 slices; TR 5.24 ms; TE 2.33 ms; flip angle 28°.

Two TMS-fMRI runs (one with ADC-fMRI and one with BOLD-fMRI) of 5 minutes and 48 seconds each were acquired in each subject. The order of TMS-fMRI runs was alternated from one subject to the next to avoid any systematic impact of the stimulation in a specific contrast. Image acquisition parameters were matched between the ADC- and BOLD-fMRI runs and were as follows: TR 1200 ms; matrix size: 72 × 72; 12 slices; 3 mm in-plane resolution; 3 mm slice thickness with 40% slice gap; in-plane GRAPPA acceleration factor 2; partial Fourier factor 6/8; simultaneous multi-slice factor 2; resulting in 216 × 216 × 50 mm^3^ partial brain coverage which comprises the M1, supplementary motor area (SMA) and primary sensory cortex (S1).

ADC-fMRI data were acquired with a custom diffusion-weighted spin-echo EPI sequence using spherical b-tensor encoding with compensation for concomitant gradient effects [32, 33] to enable isotropic functional sensitivity across fiber orientations [7]. This diffusion encoding imparts sensitivity to random displacements in all directions of space in a single shot, as opposed to linear encoding which encodes displacements along a single spatial direction at a time. The ADC-fMRI run consisted of two b = 0 volumes, followed by alternating volumes at b = 200 and 1000 s mm^−2^ (Figure 1C) at TE = 83 ms. A total of 290 diffusion-weighted volumes were acquired per run, resulting in 144 ADC timepoints with a temporal resolution of 2.4 s. The individual b-value timeseries (b200-dfMRI and b1000-dfMRI) similarly comprised 144 timepoints with the same temporal resolution. BOLD-fMRI data [34] were acquired with a multi-echo gradient-echo EPI sequence (TE 12.60 27.8 43.0 58.2 ms); flip angle 62°. A total of 290 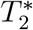-weighted volumes were acquired per run, with a temporal resolution of 1.2 s.

For both fMRI contrasts, four volumes were acquired with reverse EPI phase encoding for correction of *B*_0_ field inhomogeneity distortions. Acquisition parameters are summarised in Supplementary Table 1.

**Table 1:**
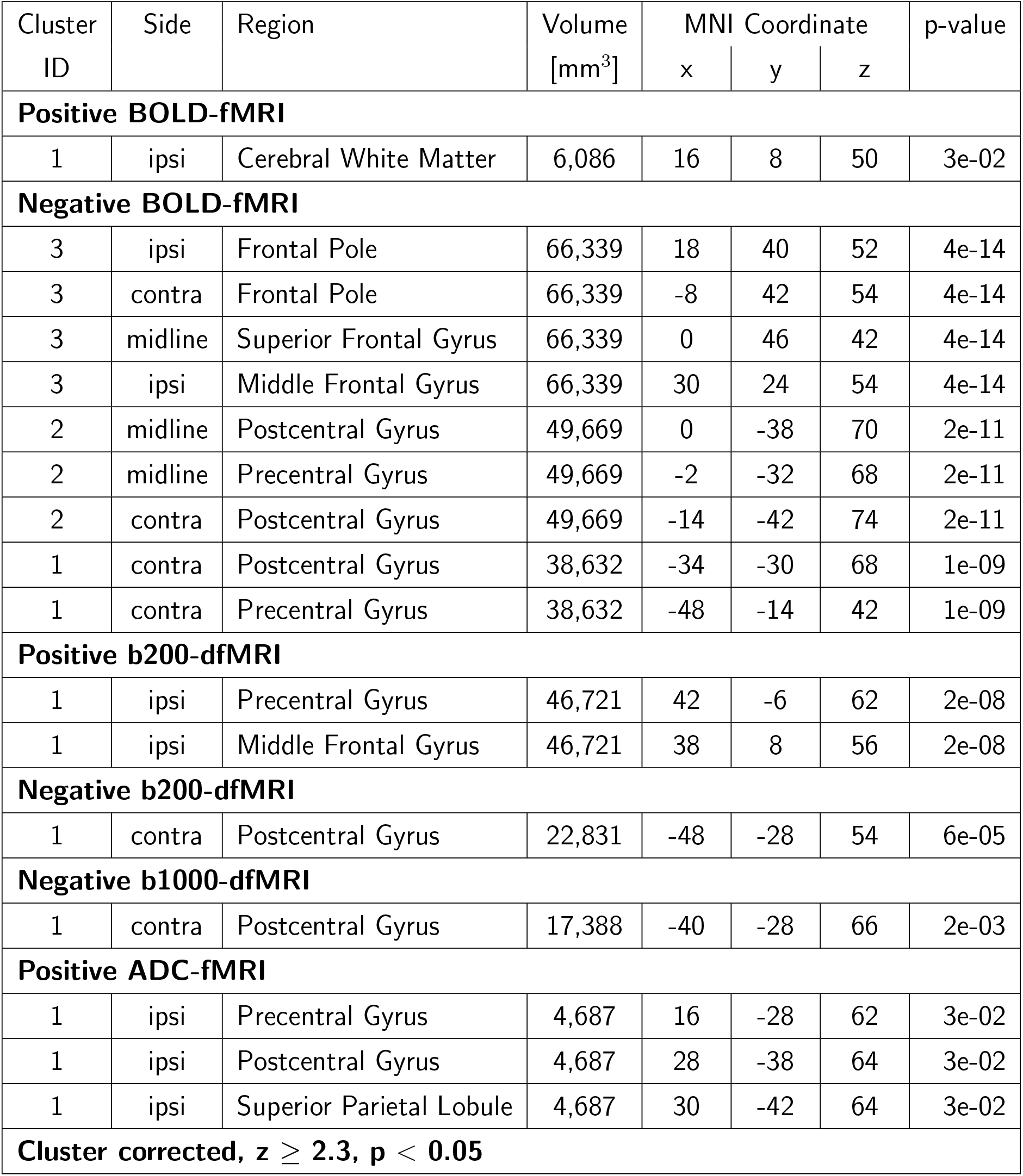
Significant clusters from group-statistics on subject-level GLM analyses. Clusters are shown for BOLD-fMRI (n=12), b200-dfMRI, b1000-dfMRI and ADC-fMRI (n=11). Sides are indicated as ipsilateral (ipsi) and contralateral (contra). Brain regions are defined according to the Harvard-Oxford atlas. Only one or two local maxima per region are shown. BOLD = blood oxygen level-dependent, ADC = apparent diffusion coefficient, dfMRI = diffusion functional MRI.

### 2.4 Transcranial Magnetic Stimulation

The TMS paradigm was conducted in a block design comprising six epochs of a baseline condition (19.2 s), followed by a TMS block (19.2 s) and a recovery period (19.2 s) (Figure 1B). During TMS blocks, 5 Hz TMS (biphasic pulse of 280 *µ*s length) was applied on the right M1 at 90% rest motor threshold (RMT), using MagVenture MagPro XP Transcranial Magnetic Stimulator. The TMS-induced current direction was anterior medial [35]. During stimulation blocks, TMS pulses were interleaved with the MRI acquisition to avoid image artifacts (Fig. 1B), as previously reported in Bestmann et al. [27, 28] and Shimomura et al. [29]. Concretely, after each trigger sent by the MRI scanner at every TR within the stimulation block, an in-house Matlab script waited 130 ms before sending a train of six pulses at 5 Hz, to allow for the MRI acquisition time (Supplementary Figure 1). This resulted in pre/post TMS pulse periods of 15/70 and 16/96 ms for ADC-fMRI and BOLD-fMRI, respectively. No TMS artifact appeared in the image, confirming that this post-TMS pulse period was sufficiently long.

### 2.5 Data Analysis

#### 2.5.1 Physiological Recordings

Physiological signals were downsampled to 100 Hz. Spurious peaks or troughs were manually clipped or replaced with the mean value of the signal to remove outliers. Bandpass filtering using Butterworth filter was applied on respiration (4^*th*^ order, 0.1-1 Hz) and cardiac (7^*th*^ order, 0.5-3 Hz) data. RETROICOR [36] was applied to extract the two first harmonics of respiration and cardiac traces, using physio_calc in AFNI.

#### 2.5.2 MRI Data

T_1_-weighted and bSSFP images were denoised with spatially adaptive non-local means filtering with ANTs [37]. Then, N4 bias field correction [38] was applied using ANTs, followed by skull-stripping with Synthstrip [39].

DfMRI data underwent a preprocessing pipeline depicted in Supplementary Figure 2, adapted from prior work [7]. Denoising was first applied to the magnitude images of the b = 200 and b = 1000 s mm^−2^ timeseries separately using NORDIC [40]. Both g-factor estimation and PCA-based denoising were performed with a 7×7×7 kernel anda step size of 1. Ringing artifacts were then mitigated in all volumes using Gibbs unringing as implemented in MRtrix3 [41, 42]. Bias field inhomogeneity was corrected using N4 bias field correction [38], with bias field extracted from the b = 0 images. Then, despiking was applied to the b = 200 and b = 1000 s mm^−2^ timeseries separately to remove outliers, using 3dDespike in AFNI. The median percentage of timepoints labelled as “big edits” was smaller than 3.5%. To correct distortion effects, susceptibility off-resonance field were estimated from the b = 0 images with forward and reversed EPI phase-encoding directions and applied across all diffusion volumes using FSL Topup [43, 44]. Motion correction was performed separately for each b-value timeseries using ANTs [45], with all volumes registered to the first b = 0 image. A brain mask was generated from the distortion-corrected b =0 images using SynthStrip [39], manually corrected in freeview from Freesurfer, and applied to exclude non-brain voxels. The functional data were sequentially registered to bSSFP 7ch, bSSFP BC, T_1_, and 2 mm MNI152 standard space, using ANTs (Supplementary Figure 2). M1, S1 and SMA from the 2 mm Harvard-Oxford cortical atlas were registered to the functional subject-space for region-of-interest (ROI) analysis. Finally, corrected b = 200 and b = 1000 timeseries were used to compute an ADC timeseries according to Equation 1. Subsequent analyses were conducted on these ADC-fMRI timeseries, as well as on the individual b-value timeseries (b200-dfMRI and b1000-dfMRI).

For BOLD-fMRI data, each echo underwent denoising, Gibbs unringing, bias field correction (bias field extracted from the first echo), and despiking procedures. The median percentage of timepoints labelled as “big edits” was smaller than 2%. Motion parameters were estimated from the first-echo timeseries, and the resulting transformations were then applied to all echoes, using ANTs. A weighted average of echoes was used to calculate an optimally combined signal, using Tedana (including denoising with TEDPCA and TEDICA) [46]. Susceptibility off-resonance field was calculated from the first echo using FSL Topup, and applied to the optimally combined data to correct distortions. The first-echo images were used to generate a brain mask, which was manually corrected and subsequently applied to remove non-brain voxels. M1, S1 and SMA from the 2 mm Harvard-Oxford atlas were registered to the functional subject-space for ROI analysis.

For quality control of functional runs, absolute displacement from the first volume and framewise displacement between volumes [47] were calculated from motion correction parameters for BOLD-fMRI, b200-dfMRI and b1000-dfMRI (Supplementary Table 2). Two epochs with framewise displacement larger than 0.2 mm were discarded in one subject, in b200-dfMRI, b1000-dfMRI, and ADC-fMRI. B200-dfMRI, b1000-dfMRI, and ADC-fMRI data from one participant were excluded because spatially varying field inhomogeneities obscured the signal. Finally, image signal-to-noise ratio (SNR) maps, defined as the average denoised signal divided by the standard deviation of the noise components (residuals), are shown in Supplementary Figure 3.

### 2.6 Statistical Analysis

The timeseries were averaged across voxels after temporal filtering (*>*0.01 Hz) and regressing out motion parameters and physiological signals from the each voxel’s timeseries. Epochs were normalised to half their baseline period (9.6 s preceding task onset) and subsequently averaged across epochs and subjects, for the left and right M1/S1 and the bilateral SMA. The power spectral density (PSD) was computed from the mean timeseries using Welch’s method implemented in SciPy [48].

Subject-level GLM analyses were performed using FSL FEAT, with temporal filtering (*>*0.01 Hz), nuisance regression (six motion parameters and the RETROICOR harmonics), and cluster correction (z ≥ 2.3, p *<* 0.05), to detect positive or negative association with the stimulation, modelled as a boxcar function. Similar GLM analyses with a boxcar function convolved with the double-gamma haemodynamic response function (HRF) can be found in the Supplementary Materials and Methods. For group analysis, the z-score maps were registered to the standard space. More specifically, the affine transformation matrices from the registration steps to T_1_ space were combined and applied as one transformation to minimise interpolation. The registration to MNI standard space and the resampling to 2 mm isotropic resolution were performed in a second step. Group-level GLMs were run with FLAME (FMRIB’s Local Analysis of Mixed Effects) stage 1 and 2, and cluster correction (z ≥ 2.3, p *<* 0.05).

We performed a two-way analysis of variance (ANOVA) on the percentage of significant voxels. For M1/S1, the two between-subject factors were the brain hemisphere (ipsilateral vs contralateral) and measured response polarity (positive vs negative association with the task). For SMA, the between-subject factor was the measured response polarity. Following the ANOVA, post-hoc pairwise comparisons were conducted using Tukey’s honestly significant difference (HSD) test to further examine differences between levels of the significant factors determined with ANOVA.

## 3 Results

### 3.1 ROI-average haemodynamic and diffusion responses

First, we investigated the ROI-average response in M1/S1 (Figure 2) and SMA (Supplementary Figure 4).

**Figure 2.**
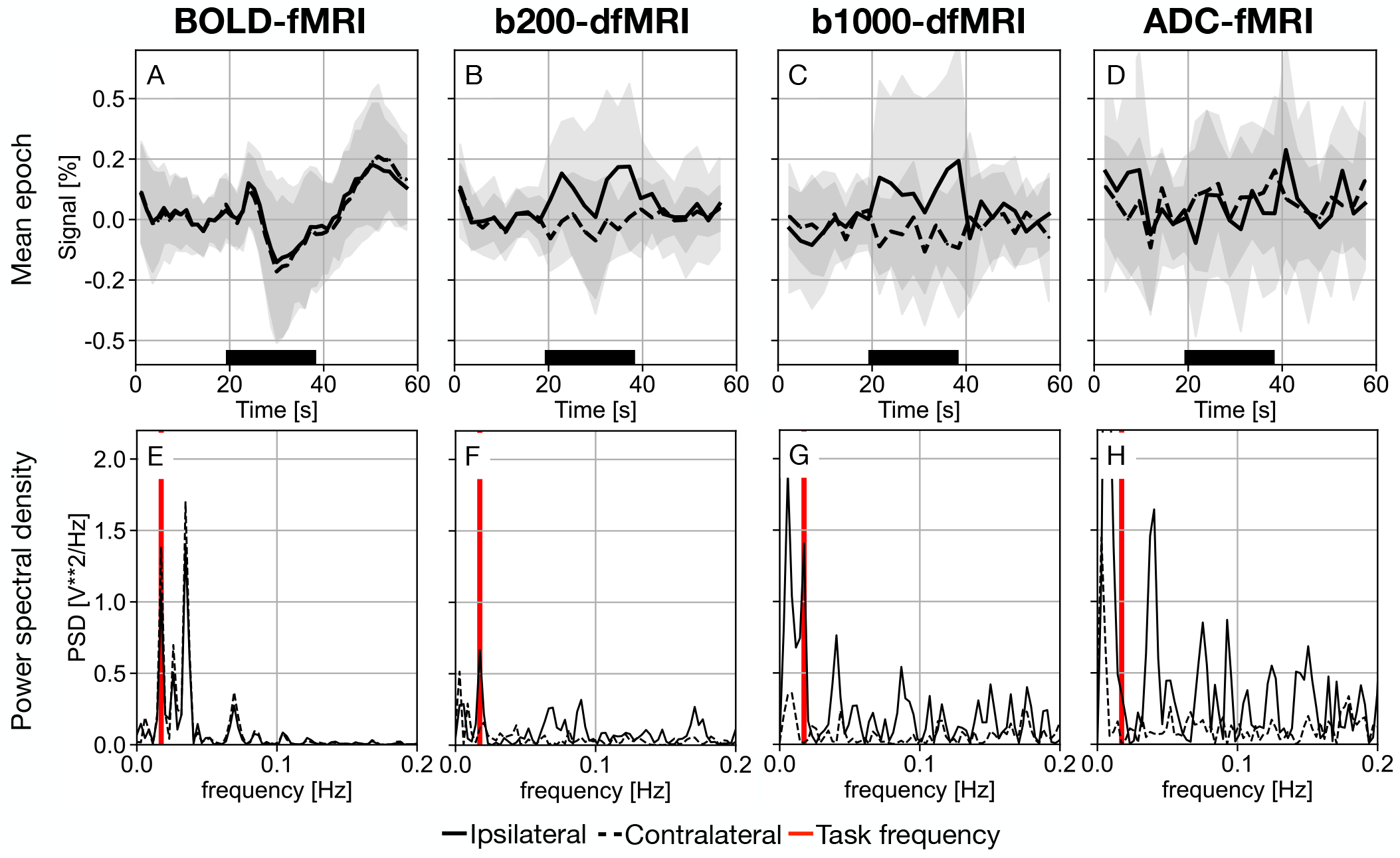
ROI-average response in M1/S1. Mean epoch time courses (A-D) and corresponding power spectral density (PSD) estimates (E-H) are shown for BOLD-fMRI (n=12), b200-dfMRI, b1000-dfMRI, and ADC-fMRI (n=11). The timeseries are averaged individually across all primary motor and somatosensory cortex (M1/S1) voxels, and then across subjects. The standard deviation across subjects is depicted as shaded grey areas. TMS duration is indicated by the black bars. The ipsilateral (ipsi, right) hemisphere is shown with solid lines, while the contralateral (contra, left) hemisphere is shown with dashed lines. The task frequency is shown on the PSD as a vertical red line. BOLD = blood oxygen level-dependent, ADC = apparent diffusion coefficient, dfMRI = diffusion functional MRI.

For BOLD-fMRI, the mean M1/S1 signal in both ipsilateral and contralateral hemispheres exhibited a small increase of up to ~ 0.1% approximately 5 s after task onset, followed by a decrease below baseline up to ~ −0.2% at 11 s after the task onset (Figure 2A). This biphasic behavior was already observed previously [49, 50]. A second positive peak of ~ 0.2% appeared around 10 s after task offset. This pattern was reflected in the PSD, which showed a peak at the task frequency and another peak at twice the task frequency (Figure 2E). The SMA mean response showed two positive peaks of ~ 0.2% (Supplementary Figure 4A), at approximately 5 s and 30 s after task onset, with a corresponding PSD peak at twice the task frequency (Supplementary Figure 4E).

For b200- and b1000-dfMRI timeseries, the M1/S1 response differed between hemispheres (Figure 2B-C). The ipsilateral response increased up to ~ 0.2% during the TMS task, with a peak in the PSD at the task frequency (Figure 2F-G), whereas the contralateral response remained essentially flat. In SMA, the response was slightly below baseline (Supplementary Figure 4B-C).

For ADC-fMRI, the average M1/S1 response in both hemispheres remained flat (Figure 2D), while the SMA response appeared positive in association with the task (Supplementary Figure 4D).

### 3.2 Group-level spatial maps

We then performed group statistics on the first-level GLM analyses using a boxcar function to identify significant clusters (Figure 3). All the axial slices are displayed in Supplementary Figure (boxcar). Positive and negative clusters identified in the group-level spatial maps are reported in Table 1. Results of GLM analyses using a boxcar function convolved with the double-gamma HRF are displayed in Supplementary Figure (HRF).

**Figure 3.**
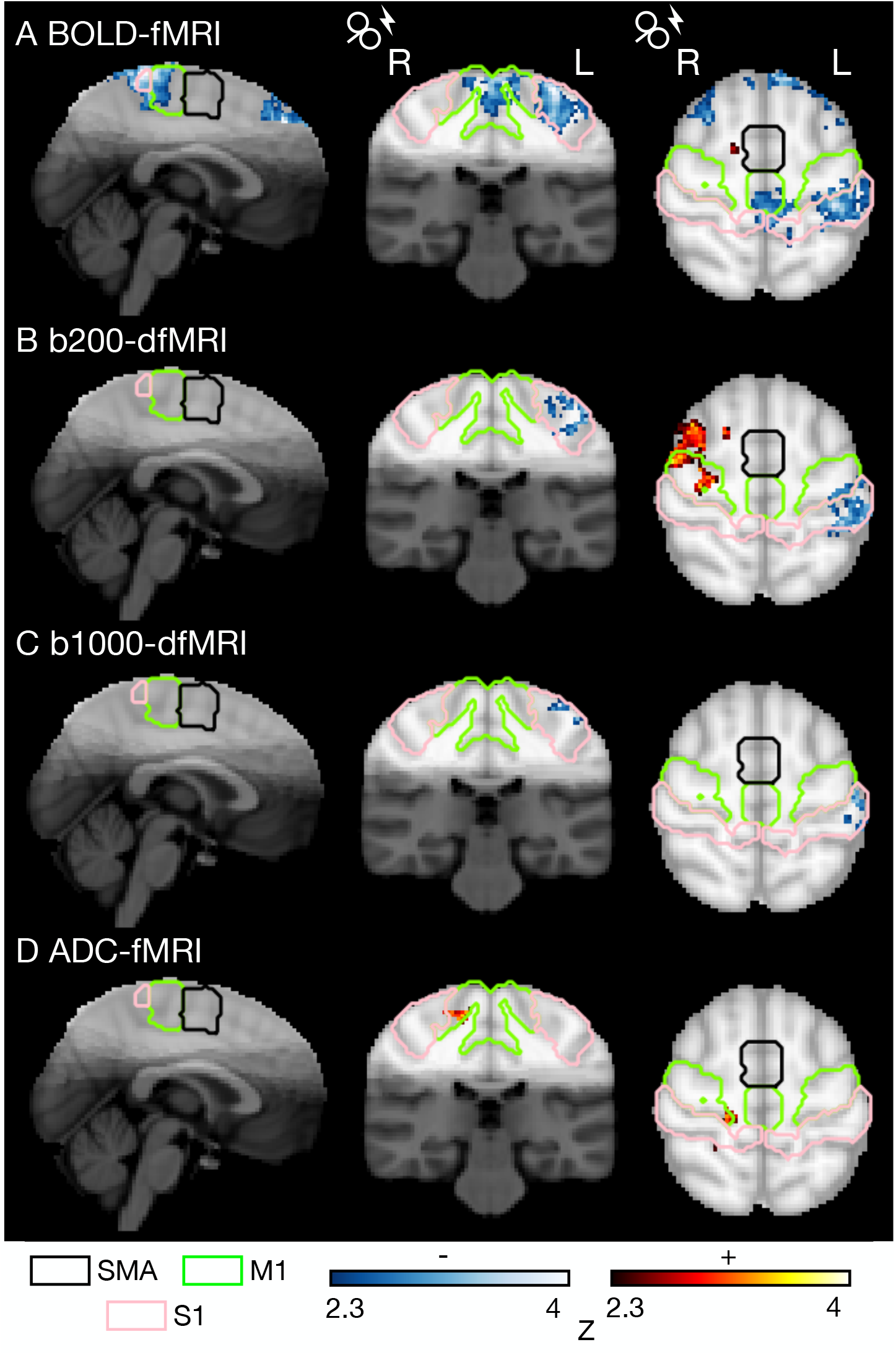
Group-level activation maps. Activation maps for (A) BOLD-fMRI (n=12), (B) b200-dfMRI, (C) b1000-dfMRI, and (D) ADC-fMRI (n=11). Clusters positively (red) and negatively (blue) correlated with the TMS paradigm are shown (general linear model; cluster-corrected; z ≥ 2.3; p *<* 0.05). The greyscale overlay represents the overlap of partial brain coverage across participants on the 2 mm MNI152 template. Anatomical ROIs from the Harvard-Oxford atlas are outlined: supplementary motor area (SMA, black), precentral gyrus/primary motor cortex (M1, green), and postcentral gyrus/primary somatosensory cortex (S1, pink). BOLD = blood oxygen level-dependent, ADC = apparent diffusion coefficient, dfMRI = diffusion functional MRI.

BOLD-fMRI yielded negative clusters in M1/S1 at the midline and contralaterally, as well as in both bilateral middle and superior frontal gyri (MFG, SFG) (Figure 3A). Additional negative BOLD clusters were detected in the bilateral frontal pole (FP). In contrast to previous work, no positive [27, 28] or negative [29] clusters were detected in the contralateral dorsal premotor cortex (PMC) or bilateral SMA.

No positive BOLD clusters were found in ipsilateral M1/S1, but a small positive cluster was observed in the ipsilateral cerebral white matter near the SMA. HRF modelling detected larger negative clusters in the middle M1/S1 and precuneus, and resulted in more spatially localised frontal clusters within the bilateral FP (Supplementary Figure 5A, HRF). Conversely, the small positive cluster in cerebral WM revealed by the boxcar response function was not captured when using the HRF.

Both dfMRI contrasts revealed a negative cluster in contralateral M1/S1 (Figure 3B-C), as reported in BOLD-fMRI. Notably, b200-dfMRI exhibited an additional positive cluster located between the ipsilateral M1 and MFG, which was absent in b1000-dfMRI and in BOLD. Similar but smaller clusters were observed with HRF modelling (Supplementary Figure 5B-C, HRF).

In ADC-fMRI, a small positive cluster emerged in ipsilateral M1/S1, extending into the superior parietal lobule (SPL), but not overlapping with the b200-dfMRI positive cluster (Figure 3D). No negative clusters survived correction for multiple comparisons. Overall, clusters detected with b1000-dfMRI and ADC-fMRI were relatively small. The mean (*±* STD) SNR within M1/S1 was 41 *±* 11 and 19 *±* 5 in b200- and b1000-dfMRI, respectively (Supplementary Table 3). In comparison, BOLD-fMRI SNR was 130 *±* 32 for TE_1_ and 71 *±* 22 for TE_4_. With HRF modelling, no cluster survived correction for multiple comparisons (Supplementary Figure 5D, HRF).

Examining individual spatial maps revealed substantial inter-subject variability (Supplementary Figure 6). The individual BOLD maps obtained using a GLM with a boxcar function convolved with the HRF yielded higher z-scores compared with the boxcar function alone (Supplementary Figure 7). The opposite effect was observed with diffusion contrasts.

### 3.3 Mean haemodynamic and diffusion responses in significant voxels

We also reported the mean responses in subject-level significant voxels, separated by polarity. Positive BOLD responses in M1/S1 (Figure 4A) and SMA (Supplementary Figure 8A) exhibited the characteristic delayed onset and gradual return to baseline, despite using a boxcar response function rather than a haemodynamic response function convolved with a boxcar. Negative BOLD responses displayed an even longer delay to onset and delay to peak [19], followed by a positive overshoot approximately 10 s after the task block ended. In SMA, a rebound was observed around 5 s after task onset. When using a boxcar function convolved with the HRF (Supplementary Figure 9A, 10A), the timeseries displayed a similar shape.

**Figure 4.**
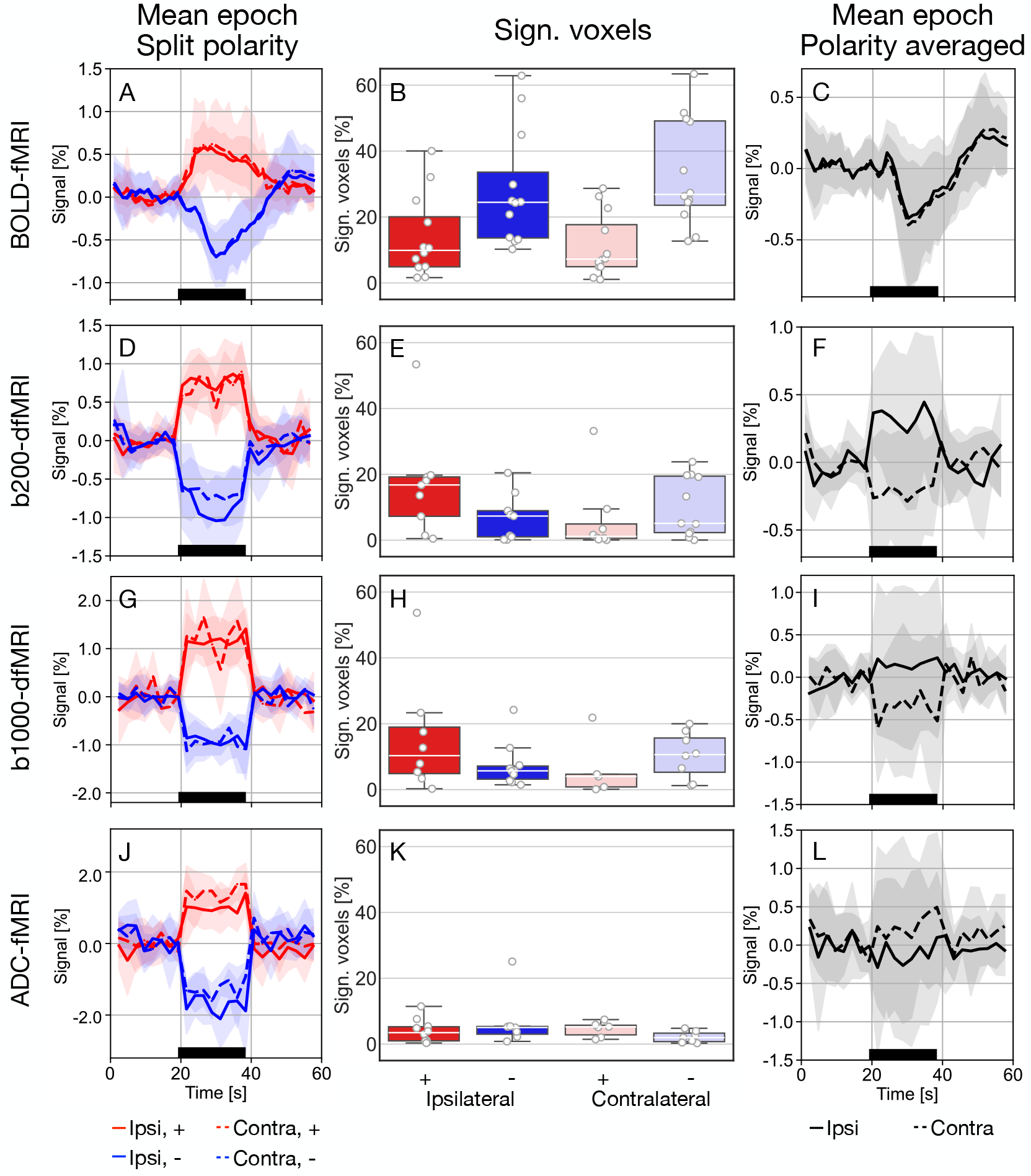
Mean response in significant voxels within M1/S1. Rows correspond to BOLD-fMRI (n=12), b200-dfMRI, b1000-dfMRI, and ADC-fMRI (n=11). Columns show mean epoch time courses in significant voxels within the primary motor and somatosensory cortex (M1/S1) split by polarity (left), corresponding percentage of significant voxels (middle), and mean epoch time courses averaged across polarities (right). Timeseries were first averaged across significant M1/S1 voxels within each subject and then across subjects. Shaded grey areas indicate the standard deviation across subjects. TMS duration is indicated by black bars. The ipsilateral (ipsi, right) hemisphere is shown with solid lines, and the contralateral (contra, left) hemisphere is shown with dashed lines. Positive (+) and negative (−) associations are depicted in red and blue, respectively. Number of subjects contributing to the averaging: n_*ipsi*,+_ = 12, 9, 7, 9, n_*ipsi*,−_ = 12, 8, 9, 7, n_*contra*,+_ = 12, 7, 5, 6, and n_*contra*,−_ = 12, 9, 7, 6, for BOLD-fMRI, b200-dfMRI, b1000-dfMRI and ADC-fMRI, respectively. BOLD = blood oxygen level-dependent, ADC = apparent diffusion coefficient.

In contrast, dfMRI and ADC-fMRI average responses (Figure 4D, G, J and Supplementary Figure 8D, G, J) displayed a sharp onset and return to baseline at the task boundaries, without any notably different shape. In this case, HRF modelling selected different voxels, leading to more gradual signal increases and decreases at the task boundaries (Supplementary Figure 9D, G, J, 10D, G, J).

Unexpectedly, ipsi- and contralateral response amplitudes in M1/S1 closely mirrored each other for all functional contrasts. This is contrary to the expectation that contralateral responses in BOLD and dfMRI would be predominantly negative and would peak at a higher amplitude than hypothesised predominantly positive ipsilateral responses, due to the nature of the subthreshold TMS. The exceptions were the ipsilateral negative responses in b200-dfMRI and ADC-fMRI, which were lower than the contralateral response.

However, examining the percentage of significant voxels within M1/S1 (Figure 4, middle) revealed, as expected, a higher proportion of negative BOLD-fMRI and b1000-dfMRI responses in contralateral M1/S1. Negative BOLD responses (Figure 4B) were more prevalent than positive responses in both hemispheres, whereas dfMRI showed a trend towards more positive responses ipsilaterally and more negative responses contralaterally (Figure 4E, H), better aligned with expectations from the paradigm design. In the SMA, significant voxels were similarly distributed across positive and negative associations, with a tendency towards more positive BOLD, and more negative dfMRI association (Supplementary Figure 8, middle). Two-way ANOVA indicated a significant effect of the response polarity in BOLD-fMRI M1/S1 (F(1, 44) = 19, p = 2e-4, Bonferroni corrected, *η*_2_ = 0.31). Post-hoc Tukey’s test showed that there are significantly more negative than positive clusters (p = 6e-5). All the other tests were non-significant. When using HRF modelling, the percentage of significant negative BOLD voxels increased, while the percentage of significant positive voxels stayed comparable (Supplementary Figure 9B). For b200-dfMRI, the percentage decreased in ipsilateral and increased in contralateral, whereas everything decreased in b1000-dfMRI and in ADC-fMRI (Supplementary Figure 9E, H, K).

We then disregarded the response polarity and averaged the signal across all significant voxels in M1/S1 (Figure 4, right). In BOLD-fMRI, the contralateral negative peak was marginally smaller than its ipsilateral counterpart (Δ = −0.05%), whereas the contralateral post-TMS overshoot was marginally greater than its ipsilateral counterpart (Δ = 0.04%) (Figure 4C). In b200-dfMRI, there was a clear distinction between the ipsilateral positive responses and the contralateral negative responses (Figure 4F). Similarly, b1000-dfMRI revealed negative contralateral responses, accompanied by less pronounced ipsilateral positive responses (Figure 4I). For ADC-fMRI, the contralateral responses were positively associated with the TMS task, whereas the ipsilateral responses were not (Figure 4L). When using HRF instead of boxcar modelling, average significant BOLD-fMRI responses remained unchanged (Supplementary Figure 9C). In b200-dfMRI (Supplementary Figure 9F), the contralateral positive and negative responses that were captured with HRF modelling cancelled each other and the average significant response was flat, as opposed to the negative contralateral average response found with boxcar modelling. In b1000-dfMRI, the average significant response captured with HRF modelling was more delayed than the one captured with boxcar modelling (Supplementary Figure 9F, I).

In SMA (Supplementary Figure 8, right), the responses were similar to the mean response in the whole ROI (Supplementary Figure 4), but the negative peaks in b1000-dfMRI and the positive peak in ADC-fMRI were more marked (Supplementary Figure 8I, L). When HRF modelling was applied, the average significant response remained unchanged in BOLD-fMRI (Supplementary Figure 10C). Conversely, the amplitudes of the average response were heavily attenuated in b1000-dfMRI and ADC-fMRI (Supplementary Figure 10F, I). In b200-dfMRI, the significant positive and negative responses captured with boxcar modelling cancelled each other and the average response appeared flat (Supplementary Figure 8F). When using HRF modelling, the average significant response was negative (Supplementary Figure

To investigate the average response of all contrasts within clusters identified in Figure 3, masks were defined for the contralateral negative BOLD-fMRI cluster, the ipsilateral positive b200-dfMRI cluster, and the ipsilateral positive ADC-fMRI cluster (Figure 5). Group-average responses for all contrasts were then computed within each mask. Within the contralateral negative BOLD cluster (Figure 5A), the dfMRI responses showed a slight negative association with the TMS task, while the ADC-fMRI response showed a positive trend. Within the ipsilateral positive b200-dfMRI cluster (Figure 5B), the dfMRI and ADC-fMRI average responses showed a positive association with the task. The BOLD-fMRI average response showed an initial positive response to the task that rapidly returned to baseline before the task ended. Within the ipsilateral positive ADC-fMRI cluster (Figure 5C), the dfMRI average responses showed a positive association with the task, while the BOLD-fMRI average response showed a negative and more delayed trend.

**Figure 5.**
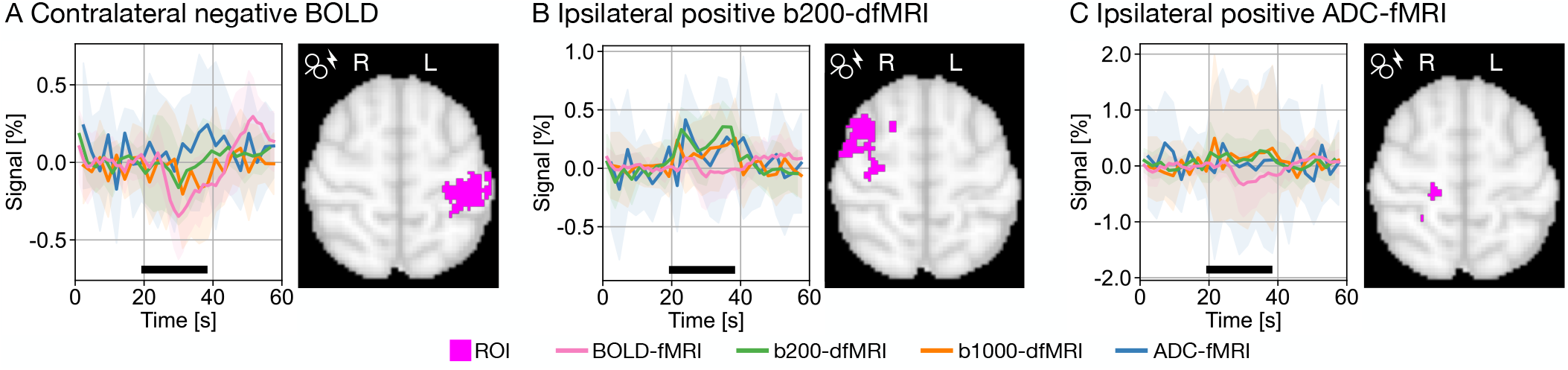
Comparison between contrasts in significant voxels. Responses are shown for BOLD-fMRI (n=12), b200-dfMRI, b1000-dfMRI and ADC-fMRI (n=11). (A) ROI corresponding to the contralateral negative BOLD-fMRI cluster shown in M1/S1 in Figure 3A. (B) ROI corresponding to the ipsilateral positive b200-dfMRI cluster shown in M1/MFG in Figure 3B. (C) ROI corresponding to the ipsilateral positive ADC-fMRI cluster shown in M1/S1 in Figure 3D. ROIs are shown as pink clusters on the 2 mm MNI152 template. Shaded areas indicate the standard deviation across subjects.

## 4 Discussion

In this study, we applied subthreshold TMS at 5 Hz on the right M1 while simultaneously acquiring fMRI data, following the design of Shimomura et al. [29]. This paradigm has been shown to elicit contralateral negative BOLD in M1/S1. We used this approach to investigate the dfMRI and ADC-fMRI counterpart of negative BOLD.

We successfully replicated all three BOLD activation clusters reported in Shimomura et al. [29] that fell within our imaging slab (negative contralateral M1/S1 and bilateral MFG). All other BOLD-fMRI clusters observed in our study were not previously reported. These include negative clusters in the bilateral FP and in the SFG along the midline, and a positive cluster in cerebral white matter adjacent to the SMA. Despite these differences, the presence of negative clusters distal to the stimulation site confirmed that subthreshold TMS effectively modulated neural circuitry without engaging corticospinal output or eliciting muscle twitches, thereby underscoring the reliability of the experimental setup.

Because BOLD arises from neurovascular coupling, its link to neuronal activity remains indirect. Diffusion fMRI offers complementary insights and may improve our understanding of the mechanisms underlying negative BOLD responses. In particular, ADC-fMRI is sensitive to excitatory neuronal activity [6–10], possibly mediated by concomitant neuromorphological rearrangements [22–25]. With appropriate experimental design, ADC-fMRI is largely independent from vascular contribution [20]. Conversely, dfMRI time courses combine T_2_-weighting and diffusion-weighting, thereby partially reflecting BOLD-like contrast while reducing direct contributions from intra-vascular signals (via diffusion-weighting) and sensitivity to macrovasculature (via spin-echo acquisition) [51, 52]. In addition, as opposed to gradient-echo BOLD, dfMRI spin-echo acquisition is not sensitive to static dephasing [51, 52].

We reported contralateral negative b200- and b1000-dfMRI clusters in M1/S1. The spatial correspondence between BOLD and dfMRI maps contralaterally demonstrates the applicability of the experimental setup beyond conventional BOLD contrast. Similarly, Pérot et al. [53] reported b200-dfMRI negative clusters in areas where high frequency visual stimulation induced negative BOLD clusters in rats. They did not find any significant ADC-fMRI response in these areas. Likewise, in our current study, no contralateral ADC-fMRI clusters survived statistical thresholding, which may indicate an inherent insensitivity of ADC functional contrast to the mechanisms driving negative BOLD, or insufficient SNR to detect microstructure-related changes of very small amplitude.

### 4.1 Ipsilateral effects of subthreshold TMS-fMRI

Consistent with previous reports, no ipsilateral M1/S1 positive BOLD cluster was observed during subthreshold TMS [27–29]. In contrast, positive BOLD clusters in ipsilateral M1/S1 were previously reported in suprathreshold TMS-fMRI paradigms (110% RMT, 3-4 Hz, 10 s ON [27, 28]). Indeed, TMS generates a phasic electric field in the underlying cortical tissue that depolarises excitable neural elements, most likely myelinated axons [54]. When the membrane potential exceeds the threshold membrane potential, it triggers an action potential that, through activation of corticospinal pyramidal neurons, can produce a muscle twitch. This neuronal excitatory activity is translated into a positive BOLD response via the haemodynamic response, and would be reflected as an ADC decrease and a b200-dfMRI and b1000-dfMRI signal increase [6–10].

In subthreshold TMS-fMRI, the absence of ipsilateral positive BOLD clusters is not trivial. Although subthreshold TMS is insufficient to elicit a muscle twitch or a positive BOLD response, it is strong enough to impact remote neuronal activity by affecting the excitatory/inhibitory balance on the contralateral side [54, 55]. This apparent paradox has been attributed to i) the lack of re-afferent feedback resulting from the absence of TMS-evoked hand movements, to ii) haemodynamic responses that remain below background physiological “noise” levels, to iii) altered haemodynamic responses not detected by conventional analysis methods, or to iv) mutual cancellation of inhibitory and excitatory effects with no haemodynamic output [27, 28]. Indeed, excitation and inhibition are not temporally distinct, but rather co-occur within local feedback loops or long-range feedforward circuits [56].

Supporting the second hypothesis, b200-dfMRI revealed a robust ipsilateral M1/PMC positive cluster that was not detected with BOLD-fMRI. This suggests that b200-dfMRI may be sensitive to subtle task-locked mechanisms not solely attributable to T_2_ effects, such as morphological changes in neurons, astrocytes or blood vessels. As the recruitment of descending elements (from cortical and motor neurons to muscle fibres), and thus MEP amplitude, scales with stimulus intensity [57, 58], an absence of muscle twitch does not mean a complete lack of neural activation or vascular response. Moreover, inhibitory interneurons appear to have a lower firing threshold than pyramidal neurons [54], making firing less energy-demanding. This could explain the negative BOLD cluster in the contralateral hemisphere in the absence of positive BOLD at the stimulation site. The absence of an ipsilateral positive cluster in b1000-dfMRI is likely attributable to its lower SNR. However, when examining the BOLD-fMRI average response within the ipsilateral positive b200-dfMRI cluster, we observed an initial increase of the BOLD amplitude that rapidly returned to baseline before the task ended. This would support the fourth hypothesis, with an early excitatory response that would be cancelled out by a later inhibitory one.

Surprisingly, rather than a negative ADC-fMRI cluster consistent with the positive b200-dfMRI cluster in M1/MFG and excitatory activity, we instead detected a positive ADC-fMRI cluster in ipsilateral M1/S1/SPL. The SPL is known for somatosensory and visuomotor integration, as well as motor learning [59]. As this cluster is not observed in BOLD, it likely reflects an ADC-specific signature rather than BOLD contamination. We could speculate that positive ADC represents the inverse of negative ADC and therefore reflects inhibition. Studies of synaptic plasticity and long-term depression have observed a link between inhibitory stimulation and morphological changes, specifically spines shrinkage, an effect reversible after excitatory stimulation [60]. While this evidence derives from long-term inhibitory processes, short-term inhibitory processes may also be accompanied by reversible microstructural rearrangements. Such spines shrinkage could contribute to the positive ADC-fMRI changes observed here, though this remains speculative and untested.

### 4.2 Contralateral effects of subthreshold TMS-fMRI

As demonstrated in previous work [27–29] and replicated in this study, subthreshold TMS applied unilaterally on M1 induces contralateral negative BOLD in M1/S1. Although negative BOLD responses are generally assumed to reflect neuronal inhibition/deactivation [5, 14, 15], their underlying mechanisms remain incompletely understood and likely do not represent a simple inverse of positive BOLD [5, 14]. Shimomura et al. [29] hypothesised that the observed contralateral negative BOLD cluster resulted from interhemispheric inhibition. Using a paired-pulse paradigm, Ferbert et al. [61] showed that EMG activity was reduced when a test pulse on the contralateral M1 followed a conditioning pulse on ipsilateral M1 after a controlled interstimulus interval. Similarly, paired-pulse transcranial eletrical stimulation reduced EMG activity and positive BOLD response in contralateral M1 [62]. Such interhemispheric inhibition likely involves excitatory transcallosal projections that excite inhibitory interneurons in contralateral M1 [54, 63, 64], or some inhibitory transcallosal projections [16, 63]. Following hyperpolarisation of postsynaptic neurons or “shunting inhibition”, corticospinal pyramidal neurons in the contralateral M1 exhibit reduced excitability [65]. In shunting inhibition, the membrane potential may remain unchanged or even become depolarised, yet neuronal excitability is still reduced [56].

Contralaterally, negative b200- and b1000-dfMRI clusters were observed in M1/S1, spatially overlapping with BOLD-fMRI negative clusters. This spatial correspondence suggests a shared underlying mechanism governing signal change in this area. In contrast, no positive ADC-fMRI cluster survived multiple comparison correction in the contralateral M1/S1. Likewise, Pérot et al. [53] did not report positive ADC-fMRI cluster in negative BOLD areas. This absence may reflect limited sensitivity of ADC-fMRI to the mechanisms underlying negative BOLD, or, alternatively, insufficient statistical power to detect more subtle neuromodulatory effects on microstructure. Should the observed negative BOLD reflect inhibition, a tentative explanation for this preferential ADC-fMRI sensitivity could be that, during inhibition and even more so during shunting inhibition, the membrane potential does not change as drastically as during action potential firing, and therefore, the associated membrane deformations may also be smaller. However, this interpretation remains speculative and requires direct experimental validation using interferometric imaging.

Regarding the potential lack of statistical power, Spencer et al. [7] detected ADC-fMRI responses to a motor task using a comparable sequence, leading to comparable SNR (Supplementary Table 3). We reiterate that b200- and b1000-dfMRI time courses have combined contributions from diffusion and BOLD-like (T_2_) effects. Thus, the detection of negative b200- and b1000-dfMRI responses in the contralateral M1/S1 does not automatically translate into an ADC change if the b200- and b1000-dfMRI changes are driven by T_2_ (microvascular) effects rather than neuromorphological effects.

### 4.3 Origin of contrasts

Notably, ADC-fMRI and dfMRI time courses exhibited sharp onsets and returns to baseline, suggesting either a diffusion-weighted contribution rather than simply mirroring the slow haemodynamic response, or, alternatively, a preferential sensitivity of spin-echo acquisition to earlier microvascular changes, as previously observed for positive BOLD [66].

While earlier responses in spin-echo vs gradient-echo positive BOLD have been reported, their temporal lead is on the order of ~ 1 s [66], which is insufficient to fully explain the larger timing differences observed between the BOLD-fMRI and the dfMRI/ADC-fMRI time-to-peak, on the order of 2-4 s. No equivalent timing analysis directly comparing spin-echo and gradient-echo acquisitions has been reported for negative BOLD responses.

A diffusion contribution to functional contrast could arise from activity-induced morphological deformations in neurons or astrocytes, but may also originate from vascular microstructure mechanisms, such as vasodilation or vasoconstriction. Regarding vascular microstructure changes, although arterial diameter begins to change within hundreds of milliseconds following neural activation, the time required to reach peak dilation and to return to baseline remains relatively slow, typically occurring several seconds after the task onset or offset [67]. Furthermore, if fluctuations in vascular microstructure were the source of the dfMRI and ADC-fMRI contrasts, we would expect a strong association between ADC-fMRI signals and breath-hold-induced brain-wide vasodilation. However, recent work has demonstrated that this association is weak to non-existent [20].

Using a similar sequence to the one in this study, Spencer et al. [7] reported sharp onsets in dfMRI time courses, likely reflecting sensitivity (via diffusion-weighting) to microstructure deformations synchronised with the task. However, they reported delayed offsets in dfMRI timeseries, which may arise from T_2_ contributions through cross-terms between diffusion-weighting gradients and susceptibility-induced background field gradients or vascular microstructure changes. In our study, the sharp nature of both onsets and offsets in dfMRI responses may suggest a contrast mechanism distinct from T_2_.

Overall, sensitivity to non-vascular microstructure changes appears to be the most plausible explanation for the sharp onsets and offsets observed in the ADC-fMRI and dfMRI time courses. However, this mechanism alone fails to explain the absence of ipsilateral clusters in b200-dfMRI and b1000-dfMRI at the location of the positive ADC-fMRI cluster.

It should be noted that, when HRF modelling was applied, dfMRI and ADC-fMRI responses exhibited more gradual onsets and offsets, yielding response profiles more similar to the haemodynamic response, as opposed to the sharp onsets and offsets originally observed using boxcar modelling. However, their onsets remained faster than those observed in BOLD, consistent with the findings of Spencer et al. [7]. Furthermore, fewer significant voxels were detected in dfMRI and ADC-fMRI when using HRF rather than boxcar modelling, suggesting that these contrasts are more sensitive to task-locked mechanisms rather than slow T_2_-related response.

Regarding neuromodulation induced by 5 Hz repetitive TMS, results should be interpreted with caution to avoid overstatement [68]. Although a relative consensus suggests that low frequency TMS (*<* 5 Hz) induces inhibitory effects and high frequency TMS (≥ 5 Hz) induces excitatory effects [58], the underlying mechanisms are considerably more complex [68]. The effects of TMS are influenced by stimulation amplitude, frequency, coil orientation, and the total number of pulses delivered, among other factors. For instance, Gorsler et al. [69] reported a right hand MEP increase after 1800 pulses of TMS (5 Hz, 95% RMT) on the right M1, which would suggest an interhemispheric facilitatory effect. Furthermore, the total number of pulses also appears to be a critical factor. In Peinemann et al. [70], an increase in right MEP following TMS on the left M1 (5 Hz, 90% RMT) was observed after 1800 pulses, whereas no such effect was detected after 150 pulses. Using a similar methodology, Quartarone et al. [71] reported that right MEP amplitudes were significantly increased after 900 pulses, but not after 300 or 600 pulses. In this study, participants received 96 TMS pulses per task block, leading to 576 pulses per functional run (six blocks) and 1152 pulses per scanning session (two runs). Taken together, these findings indicate that stimulation frequency and intensity alone are insufficient to account for TMS effects, highlighting the importance of pulse number as an additional determinant. To better characterise the effects of TMS, direct physiological measurements with the same TMS paradigm should be employed, such as MEP amplitude (via paired-pulse TMS [61]), deoxyhaemoglobin concentration (with non-infrared spectroscopy [72]), or regional cerebral blood flow (with PET [73]).

### 4.4 Limitations

This study presents some limitations and methodological challenges. First, delivering precise stimulation, both in terms of target location and intensity, is difficult. While EMG recordings of MEPs following single pulses help validate TMS accuracy, neuronavigation systems would provide even greater precision. Even small deviations from the intended target can lead to an overestimation of the RMT, potentially affecting stimulation efficacy and consistency across participants. Additionally, head displacement relative to the coil can introduce variability in the SNR, potentially obscuring signal variations of interest. This issue can be partially mitigated by motion tracking and real-time adaptation of TMS intensity [74, 75]. Although no such techniques were employed in the present study, the replication of previously reported results suggests that both the location and intensity of TMS were applied consistently. In addition, ADC-fMRI and BOLD-fMRI data were acquired in the same subjects during the same session, enabling direct comparison between the modalities.

Second, due to the geometry of the TMS coils, limiting head motion or vibration after the TMS coil positioning is challenging. During scanning, it is mitigated by fixing the participant’s head solidly in place with vacuum pads and an external strap. In GLM analysis, motion parameters and physiological signals were also regressed out to further reduce motion-related confounds.

Third, achieving adequate spatial and temporal resolution while maintaining sufficient SNR within the constraints imposed by the TMS coils remains challenging, although it can be improved through EPI acquisition, multiband acceleration, and partial Fourier. In the present study, the use of partial brain coverage introduced variability due to differences in slab positioning across participants. This partially limits anatomical overlap at the group level, particularly in regions near the edges of the acquired slab. Moreover, the bilateral auditory cortices, typically used as a control region for TMS, are absent from the imaging slab. Interhemispheric inhibition has also been shown to be stronger in the dominant hemisphere [65]. Although stimulation of the left M1 would therefore be expected to maximise interhemispheric effects in right-handed participants, technical constraints necessitated stimulation of the right M1 in the present study.

Finally, while TMS is a powerful tool to directly modulate neuronal activity, it represents an artificial modulation which interacts with brain states and does not fully replicate spontaneous engagement of the same network [68, 76].

In conclusion, our subthreshold TMS-fMRI paradigm broadly replicated previous BOLD findings, including negative BOLD responses in contralateral M1/S1 and at the midline, and absence of positive BOLD responses in ipsilateral M1/S1. We reported for the first time dfMRI and ADC-fMRI responses to TMS. We found contralateral negative dfMRI clusters that spatially overlapped with the BOLD-fMRI negative cluster in M1/S1, suggesting a shared underlying mechanism governing this area. However, the sharper onsets and offsets of the diffusion-weighted fMRI responses may indicate a contribution from mechanisms different from 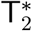 BOLD, in the form of T_2_-sensitive or diffusion-sensitive processes. No contralateral ADC clusters were observed, which suggests that the ADC-fMRI contrast either lacks a negative BOLD counterpart, or is limited by statistical power to capture very weak effects. Ipsilaterally, we reported a positive b200-dfMRI cluster in M1/PMC, potentially reflecting enhanced sensitivity to subtle ipsilateral task-locked mechanisms, below the detectability threshold of BOLD, b1000-dfMRI and ADC-fMRI contrasts. Overall, dfMRI and ADC-fMRI offer promising avenues to further disentangle neuronal and vascular contributions to complex responses of subthreshold TMS, though the precise vascular or microstructural mechanisms underlying negative BOLD, negative dfMRI, and positive ADC-fMRI remain to be fully elucidated.

## Acknowledgements

The authors thank Anne-Cerise Bernard, Loan Mattera, Nathalie Philippe, Olivier Reynaud and Rebecca Jones for their help in setting up the TMS-fMRI experiments. They thank Stefano Moia for advice regarding physiological monitoring and preprocessing. They also thank Rebecca Jones and Sarah Grosshagauer for insightful discussion. The authors acknowledge the Fondation Campus Biotech Geneva for providing expertise and resources to conduct this study.

## Funding

Swiss Secretariat for Research and Innovation (SERI), ERC Starting Grant award ‘FIREPATH’ MB22.00032 (I.J.)

SNSF Eccellenza fellowship no. 194260 (I.J.)

Swedish Cancer Society 22 0592 JIA (F.S.)

## Author contributions

Conceptualisation: I.J., I.d.R., A.S.;

Methodology: I.J., I.d.R., A.S., R.M., V.R., J-B.P, F.S.;

Validation: I.d.R.;

Formal analysis: I.d.R.;

Investigation: I.J., I.d.R., R.M., V.R.;

Data curation: I.d.R.;

Writing-original draft: I.d.R.;

Writing - Review & Editing: I.d.R, A.S., R.M., V.R., J-B.P, F.S., I.J.;

Visualisation: I.d.R.;

Supervision: I.J.;

Funding acquisition: I.J.

## Data and materials availability

The data will be available upon publication.

