## Supplementary Material for "Characterising the diffusion functional signature of negative BOLD with interleaved TMS-fMRI in the human brain"

Inès de Riedmatten *et al.*

**This PDF file includes:**

Figs. S1 to S10

Tables S1 to S2

Supplementary Table 1: **Acquisition parameters**. bSSFP BC = body coil, bSSFP 7ch = 2 x 7 channels MRI receive coils.

| Parameters | bSSFP BC | bSSFP 7ch | MPRAGE | dfMRI | BOLD |
|---|---|---|---|---|---|
| TE [ms] | 2.33 | 2.33 | 2.98 | 83 | 12.6, 27.8, 43.0, 58.2 |
| TR [s] | 5.24e-3 | 5.24e-3 | 2.3 | 1.2 | 1.2 |
| Temporal resolution [s] | - | - | - | 2.4 | 1.2 |
| Matrix | 96x96 | 96x96 | 92x240 | 72x72 | 72x72 |
| Slices | 128 | 128 | 256 | 12 | 12 |
| Resolution [$mm^3$] | 2x2x2 | 2x2x2 | 1x1x1 | 3x3x3 | 3x3x3 |
| Slice gap [%] | - | - | - | 40 | 40 |
| Flip angle [°] | 28 | 28 | 9 | 90 | 62 |
| Multiband - Grappa | - | - | 2- - | 2-2 | 2-2 |
| Partial Fourier | 1 | 1 | 1 | 6/8 | 6/8 |

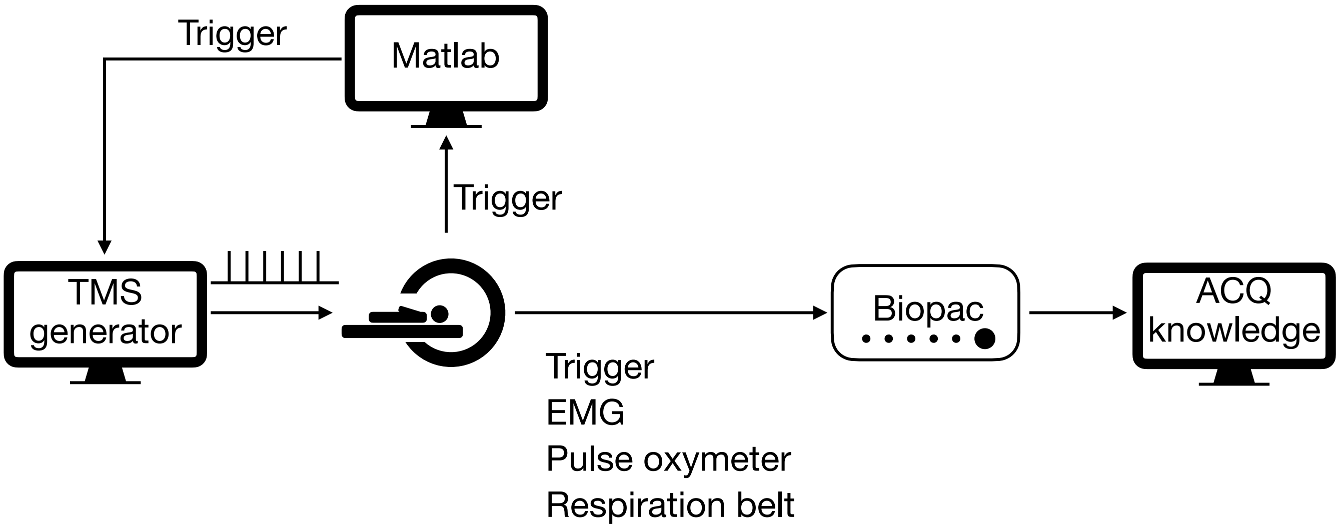


Supplementary Figure 1: **Data acquisition.** The TMS pulses are precisely timed between EPI readouts. The MRI scanner sends a trigger for each acquired volume to a MATLAB script. During task blocks, the script forwards the volume trigger to the TMS generator, which delivers a train of six pulses at 5 Hz. TMS triggers, EMG, pulse oximeter, and respiration belt signals are routed to a Biopac unit and recorded using the ACQknowledge software.

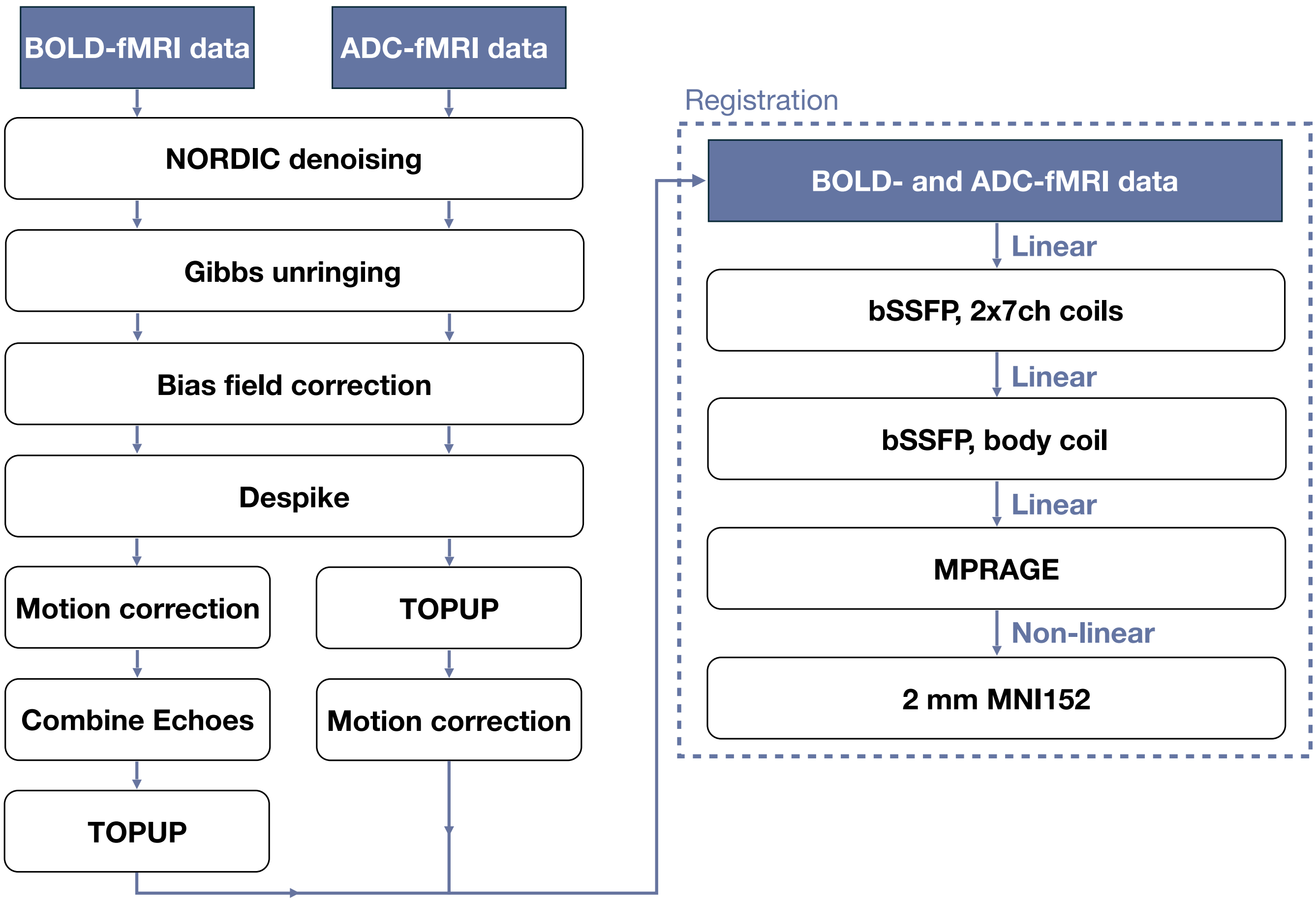


Supplementary Figure 2: **Preprocessing pipeline for BOLD-fMRI and ADC-fMRI data.**

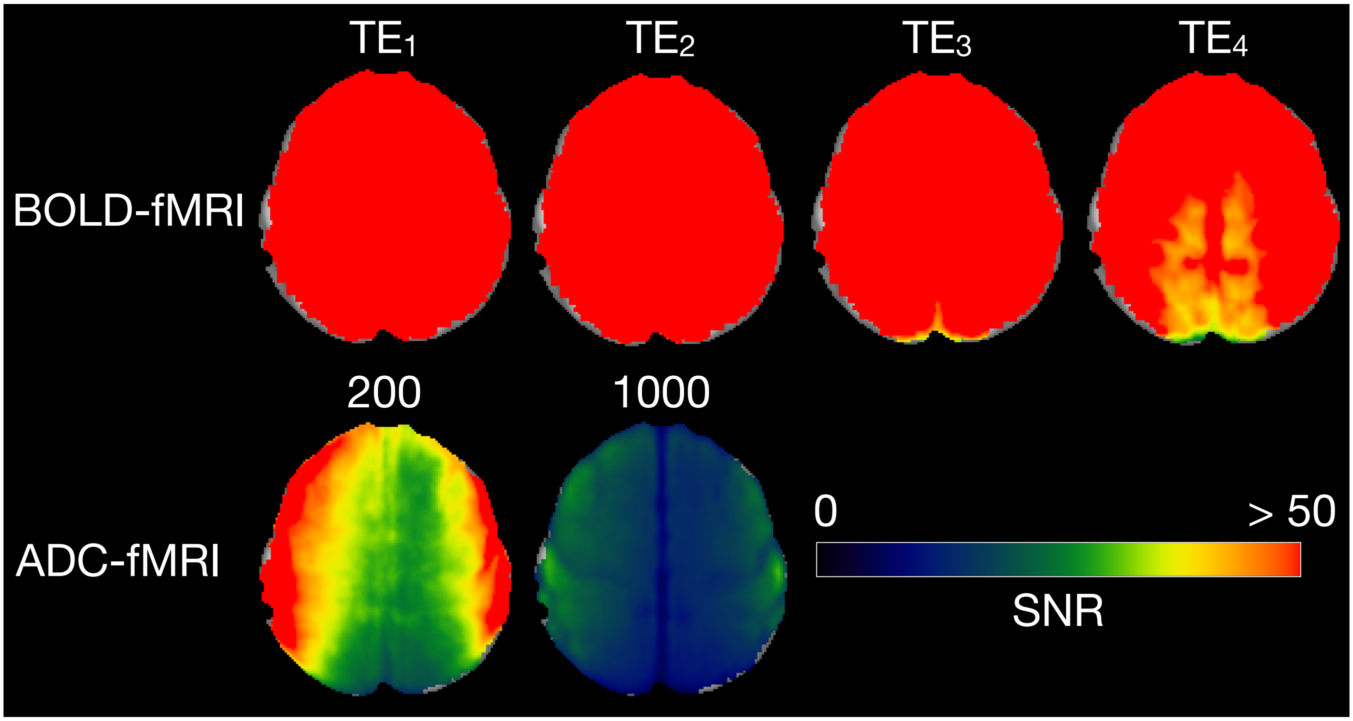


Supplementary Figure 3: **Group-mean image SNR maps for BOLD-fMRI and ADC-fMRI data.** SNR was calculated as the mean denoised signal divided by the standard deviation of the components removed by denoising (residuals). Group-mean SNR maps are shown for the four echo times (TE) from BOLD-fMRI data, and for $b = 200$ and $b = 1000$ s mm$^{-2}$ from the ADC-fMRI acquisition for comparison. Mean SNR values within the brain are shown in Supplementary Table 3. Note that image SNR values cannot be calculated for ADC-fMRI, as this is derived from the constituent b-value acquisitions.

Supplementary Table 2: **Framewise displacement for BOLD-fMRI and ADC-fMRI data.** The individual framewise displacement (FD) is calculated during the motion correction steps, using ANTS. The mean and max (standard deviation) across subjects of the FD is depitcted. Note that the FD cannot be calculated for ADC-fMRI, as this is derived from the constituent b-value acquisition.

| Mean FD [mm] | BOLD-fMRI | b = 200 | b = 1000 |
|---|---|---|---|
| | 0.11 (0.04) | 0.16 (0.04) | 0.14 (0.04) |
| Max FD [mm] | | | |
| | 0.42 (0.17) | 0.69 (0.15) | 0.62 (0.19) |

Supplementary Table 3: **Average image SNR for BOLD-fMRI and ADC-fMRI data.** Mean (standard deviation) across subjects of the average SNR within the M1/S1 regions of the Harvard-Oxford atlas are shown for the four echo times (TE) from BOLD-fMRI data, and for b = 200 and b = 1000 s mm$^{-2}$ from the ADC-fMRI acquisition for comparison. The last row shows the average SNR within the M1/S1 regions, derived from the motor results in Spencer et al. [7].

| | $TE_1$ | $TE_2$ | $TE_3$ | $TE_4$ |
|---|---|---|---|---|
| BOLD-fMRI | 130 (32) | 111 (31) | 89 (27) | 71 (22) |
| | b = 200 | | b = 1000 | |
| ADC-fMRI | 41 (11) | | 19 (5) | |
| From Spencer et al. [7] | 38 (11) | | 18 (5) | |

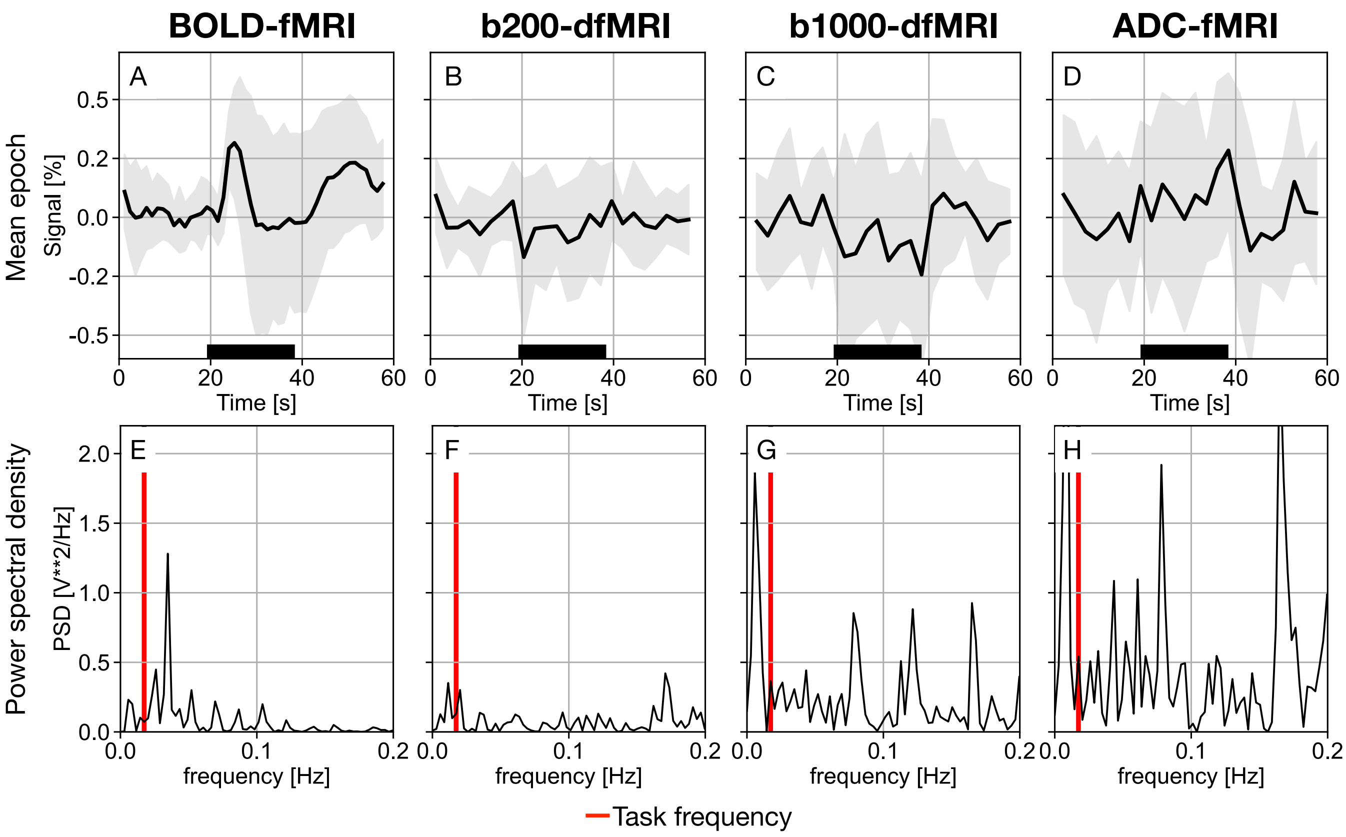


Supplementary Figure 4: **ROI-average response in SMA.** Mean epoch time courses (A-D) and corresponding power spectral density (PSD) estimates (E-H) are shown for BOLD-fMRI (n=12), b200-dfMRI, b1000-dfMRI, and ADC-fMRI (n=11). The timeseries are averaged individually across all supplementary motor area (SMA) voxels and then across subjects. The standard deviation across subjects is depicted as shaded grey areas. TMS duration is indicated by the black bars. The task frequency is shown on the PSD as a vertical red line. BOLD = blood oxygen level-dependent, ADC = apparent diffusion coefficient.

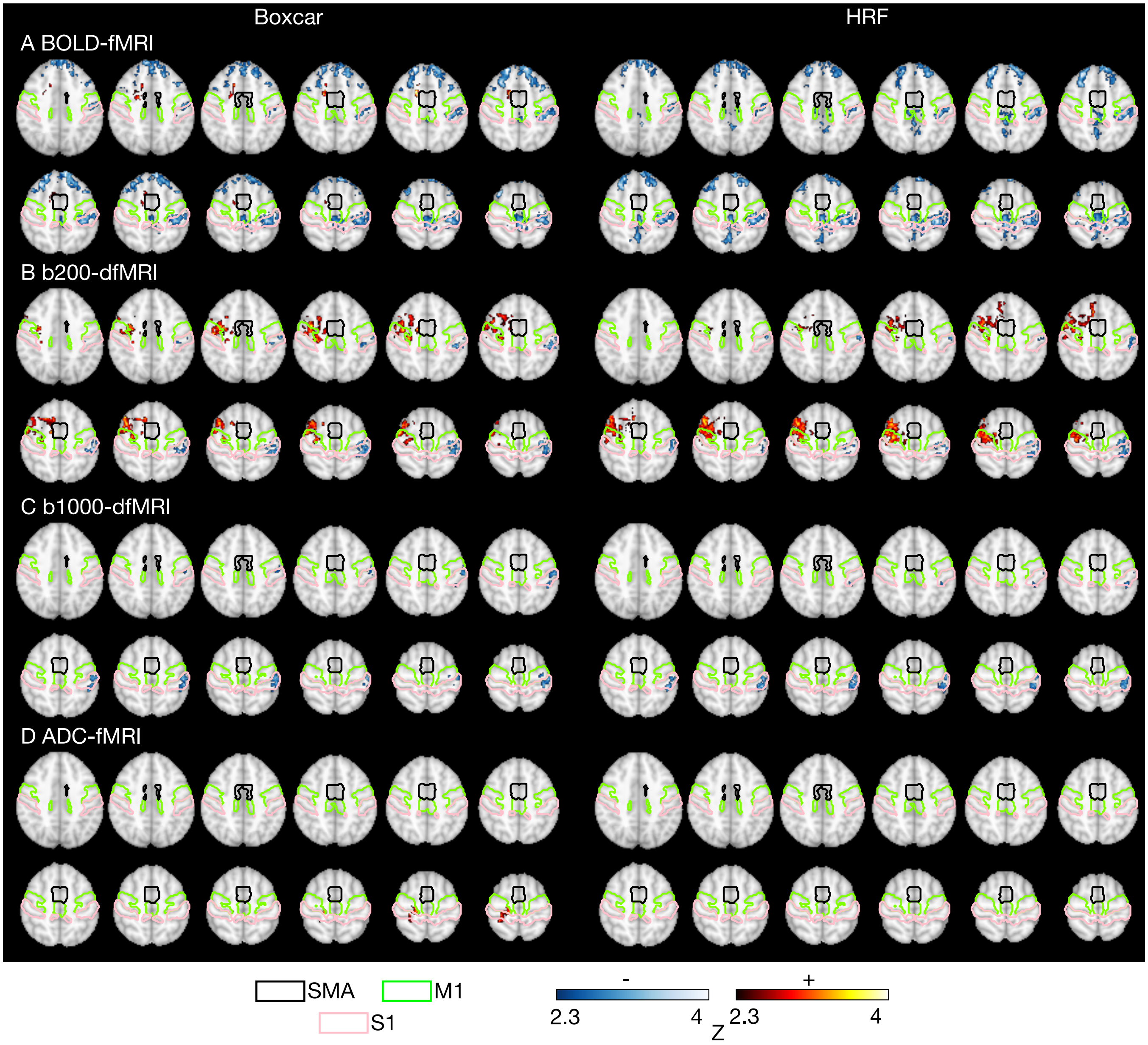


Supplementary Figure 5: **Group-level activation maps (boxcar and HRF models).** Activation maps are shown for (A) BOLD-fMRI (n=12), (B) b200-dfMRI, (C) b1000-dfMRI, and (D) ADC-fMRI (n=11). Clusters positively (red) and negatively (blue) correlated with the TMS paradigm are shown (general linear model; cluster-corrected; $z \geq 2.3$; $p < 0.05$), displayed on the 2 mm MNI152 template. The task was either modelled with a boxcar function (left column) or a boxcar function convolved with a double-gamma haemodynamic response function (HRF, right column). Anatomical ROIs from the Harvard-Oxford atlas are outlined: supplementary motor area (SMA, black), precentral gyrus/primary motor cortex (M1, green), and postcentral gyrus/primary somatosensory cortex (S1, pink). BOLD = blood oxygen level-dependent, ADC = apparent diffusion coefficient.

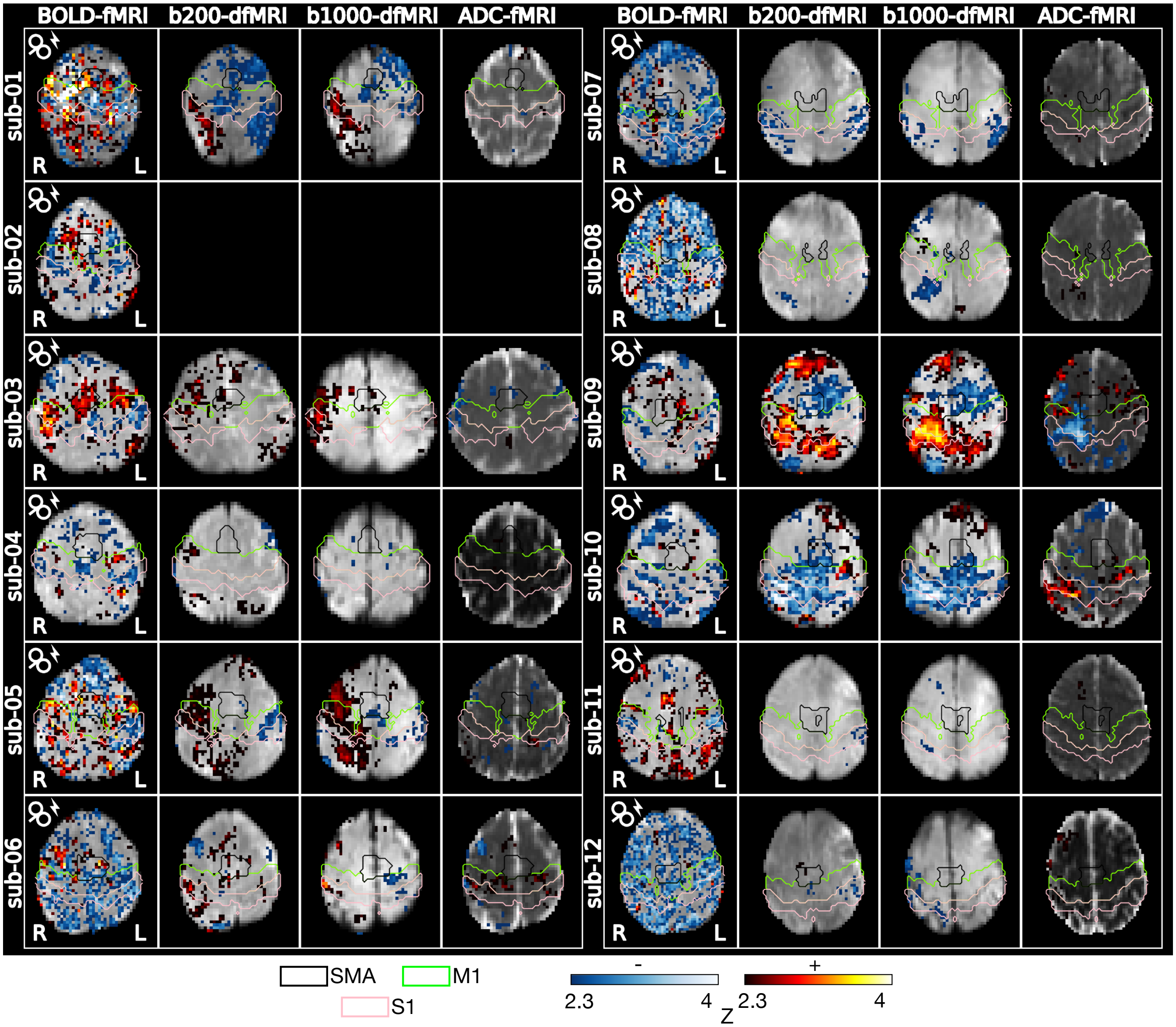


Supplementary Figure 6: **Subject-level activation maps (boxcar model)**. Activation maps are shown for BOLD-fMRI, b200-dfMRI, b1000-dfMRI and ADC-fMRI. Clusters showing positive (red) or negative (blue) association with the TMS paradigm are shown (general linear model; cluster-corrected; $z \geq 2.3$; $p < 0.05$). Anatomical ROIs from the Harvard-Oxford atlas are outlined: supplementary motor area (SMA, black), precentral gyrus/primary motor cortex (M1, green), and postcentral gyrus/primary somatosensory cortex (S1, pink). BOLD = blood oxygen level-dependent, ADC = apparent diffusion coefficient. The dfMRI run of sub-02 was discarded due to signal dropout.

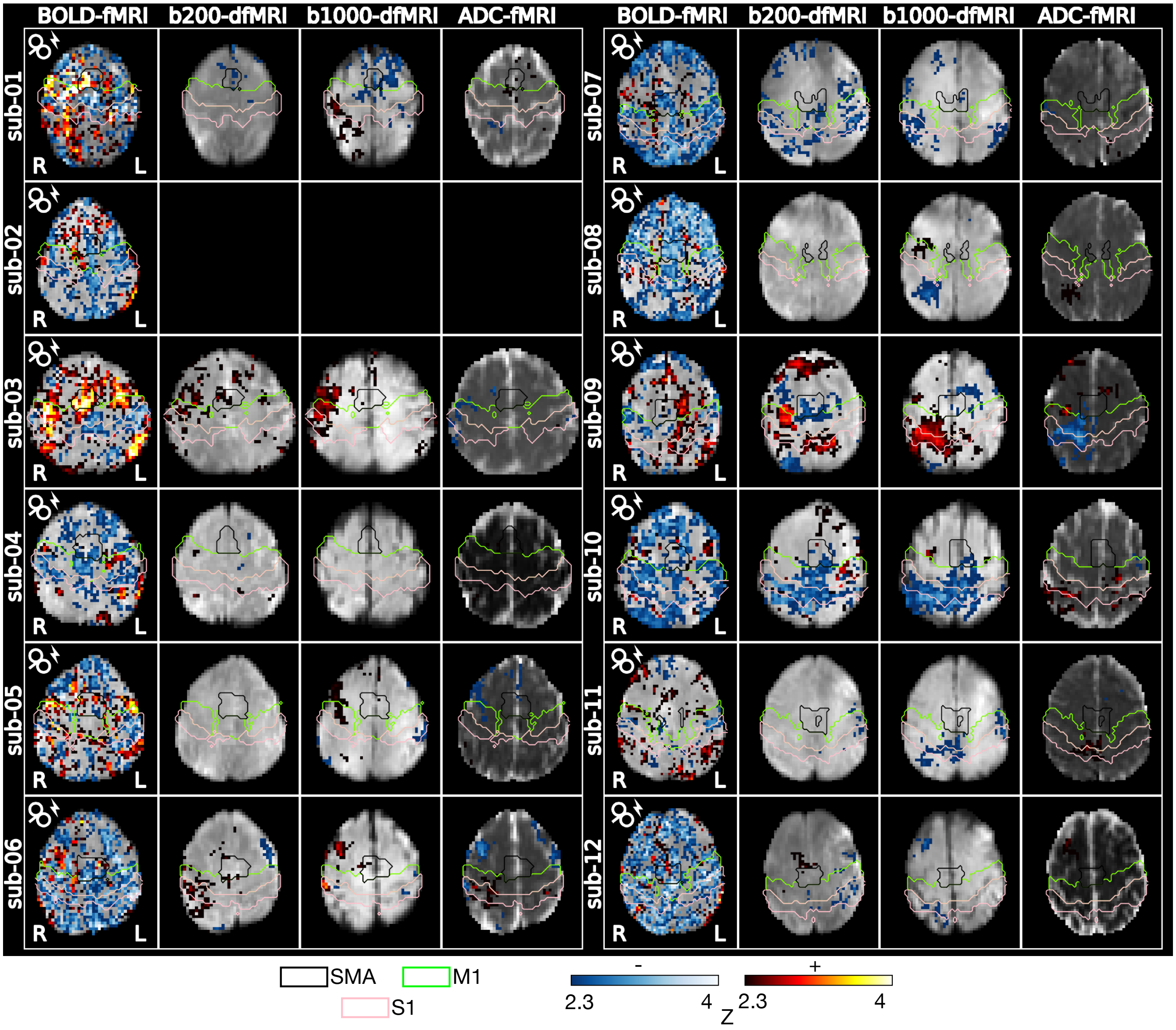


Supplementary Figure 7: **Subject-level activation maps (HRF model)**. Activation maps are shown for BOLD-fMRI, b200-dfMRI, b1000-dfMRI and ADC-fMRI. Clusters showing positive (red) or negative (blue) association with the TMS paradigm are shown (general linear model; cluster-corrected; $z \geq 2.3$; $p < 0.05$). Anatomical ROIs from the Harvard-Oxford atlas are outlined: supplementary motor area (SMA, black), precentral gyrus/primary motor cortex (M1, green), and postcentral gyrus/primary somatosensory cortex (S1, pink). BOLD = blood oxygen level-dependent, ADC = apparent diffusion coefficient. The dfMRI run of sub-02 was discarded due to signal dropout.

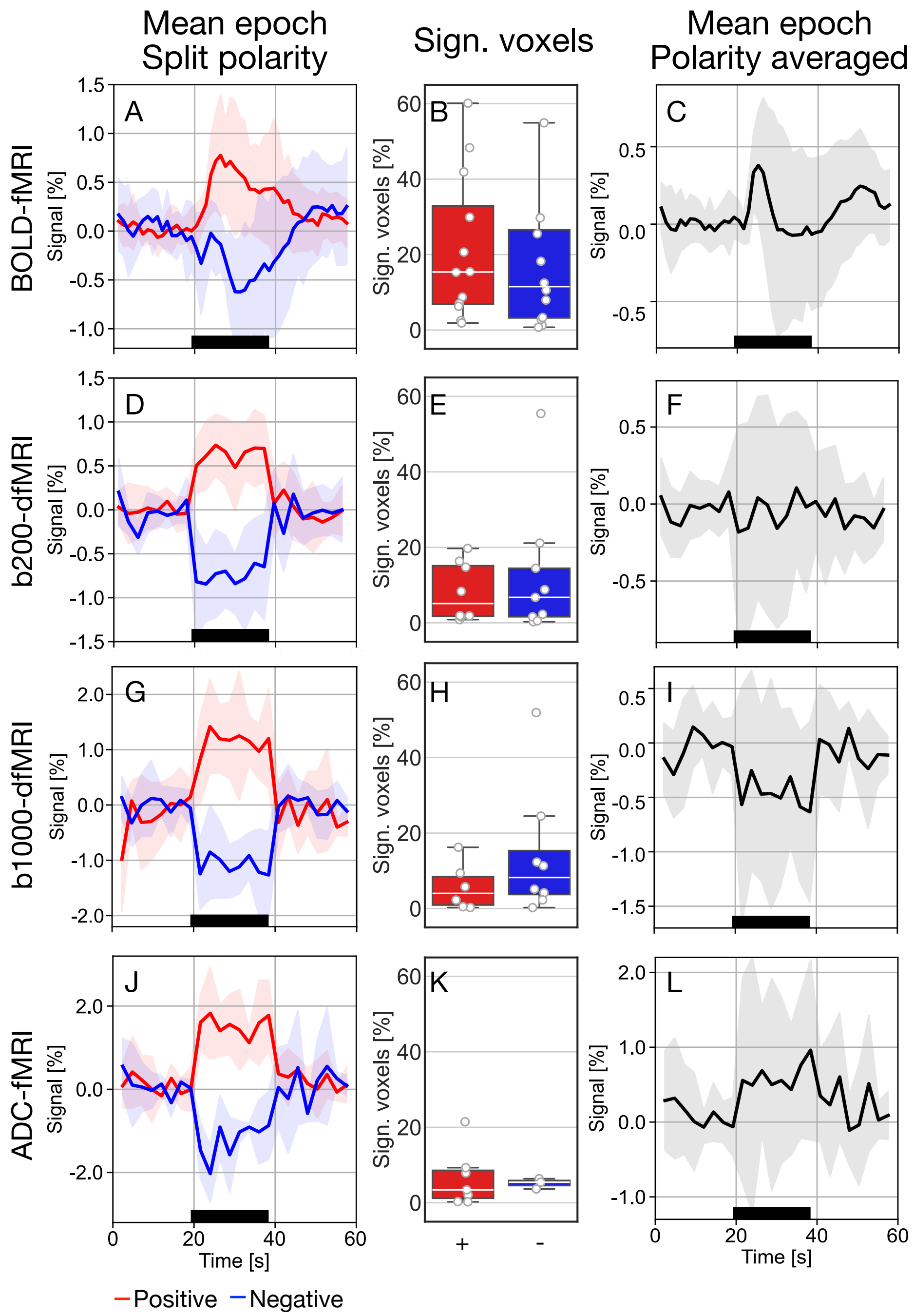


Supplementary Figure 8: **Mean response in significant voxels within SMA (boxcar model)**. Rows correspond to the BOLD-fMRI (n=12), b200-dfMRI, b1000-dfMRI, and ADC-fMRI (n=11) contrasts. Columns show mean epoch time courses in significant voxels within the supplementary motor area (SMA) split by polarity (left), corresponding percentage of significant voxels (middle), and mean epoch time courses averaged across polarities (right). Timeseries were first averaged across significant SMA voxels within each subject and then across subjects. Shaded grey areas indicate the standard deviation across subjects. TMS duration is indicated by black bars. Positive (+) and negative (-) associations are depicted in red and blue, respectively. Number of subjects contributing to the averaging: $n_{positive}$ = 12, 8, 5, 5 and $n_{negative}$ = 12, 7, 7, 3, for BOLD-fMRI, b200-dfMRI, b1000-dfMRI and ADC-fMRI, respectively. BOLD = blood oxygen level-dependent, ADC = apparent diffusion coefficient.

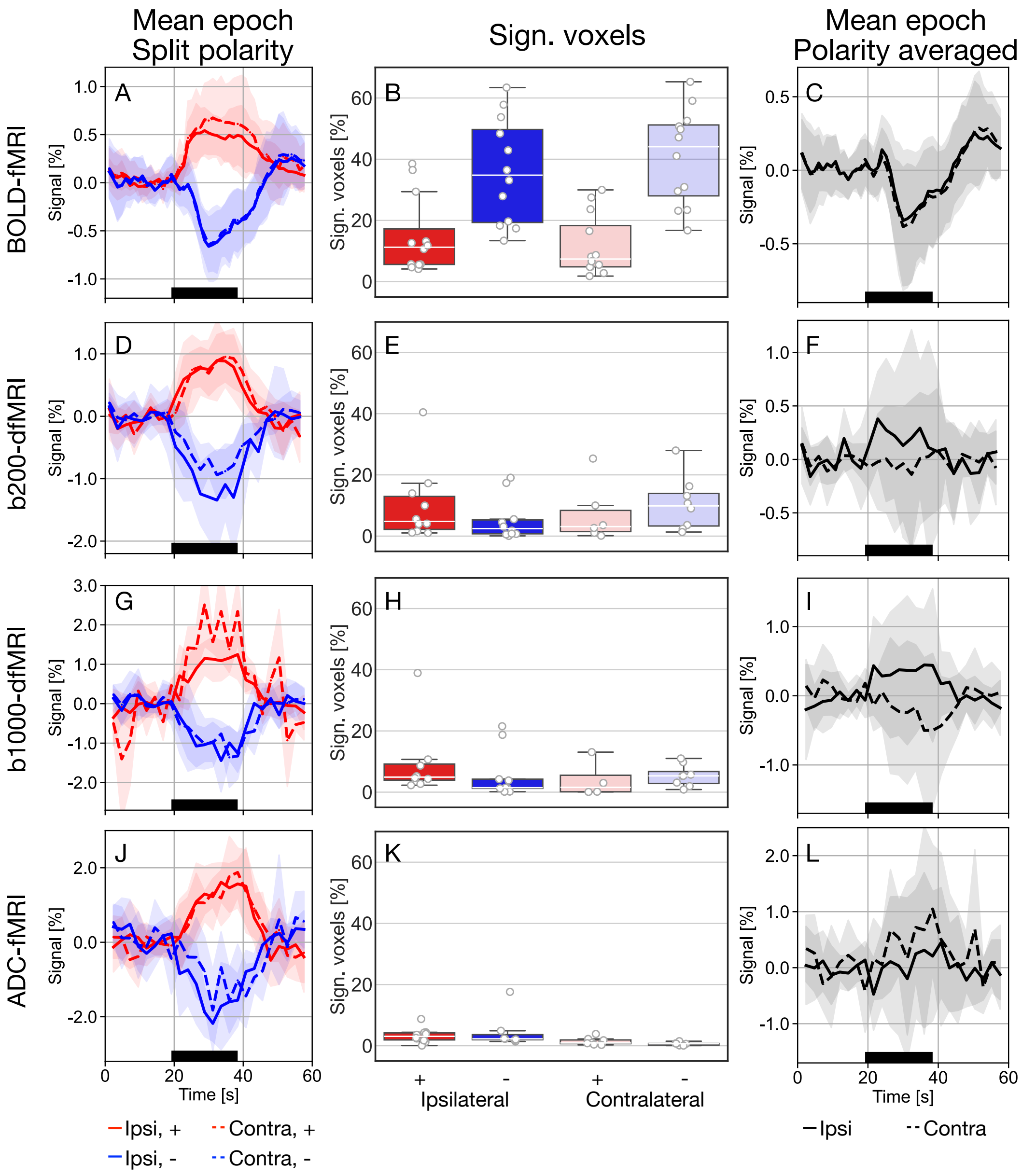


Supplementary Figure 9: **Mean response in significant voxels within M1/S1 (HRF model)**. Rows correspond to the BOLD-fMRI (n=12), b200-dfMRI, b1000-dfMRI, and ADC-fMRI (n=11) contrasts. Columns show mean epoch time courses in significant voxels within the primary motor and somatosensory cortex (M1/S1) split by polarity (left), corresponding percentage of significant voxels (middle), and mean epoch time courses averaged across polarities (right). Timeseries were first averaged across significant M1/S1 voxels within each subject and then across subjects. Shaded grey areas indicate the standard deviation across subjects. TMS duration is indicated by black bars. The ipsilateral (ipsi, right) hemisphere is shown with solid lines, and the contralateral (contra, left) hemisphere is shown with dashed lines. Positive (+) and negative (-) associations are depicted in red and blue, respectively. Number of subjects contributing to the averaging: $n_{ipsi,+}$ = 12, 9, 8, 8, $n_{ipsi,-}$ = 12, 8, 8, 6, $n_{contra,+}$ = 12, 6, 3, 7, and $n_{contra,-}$ = 12, 7, 7, 5, for BOLD-fMRI, b200-dfMRI, b1000-dfMRI and ADC-fMRI, respectively. BOLD = blood oxygen level-dependent, ADC = apparent diffusion coefficient.

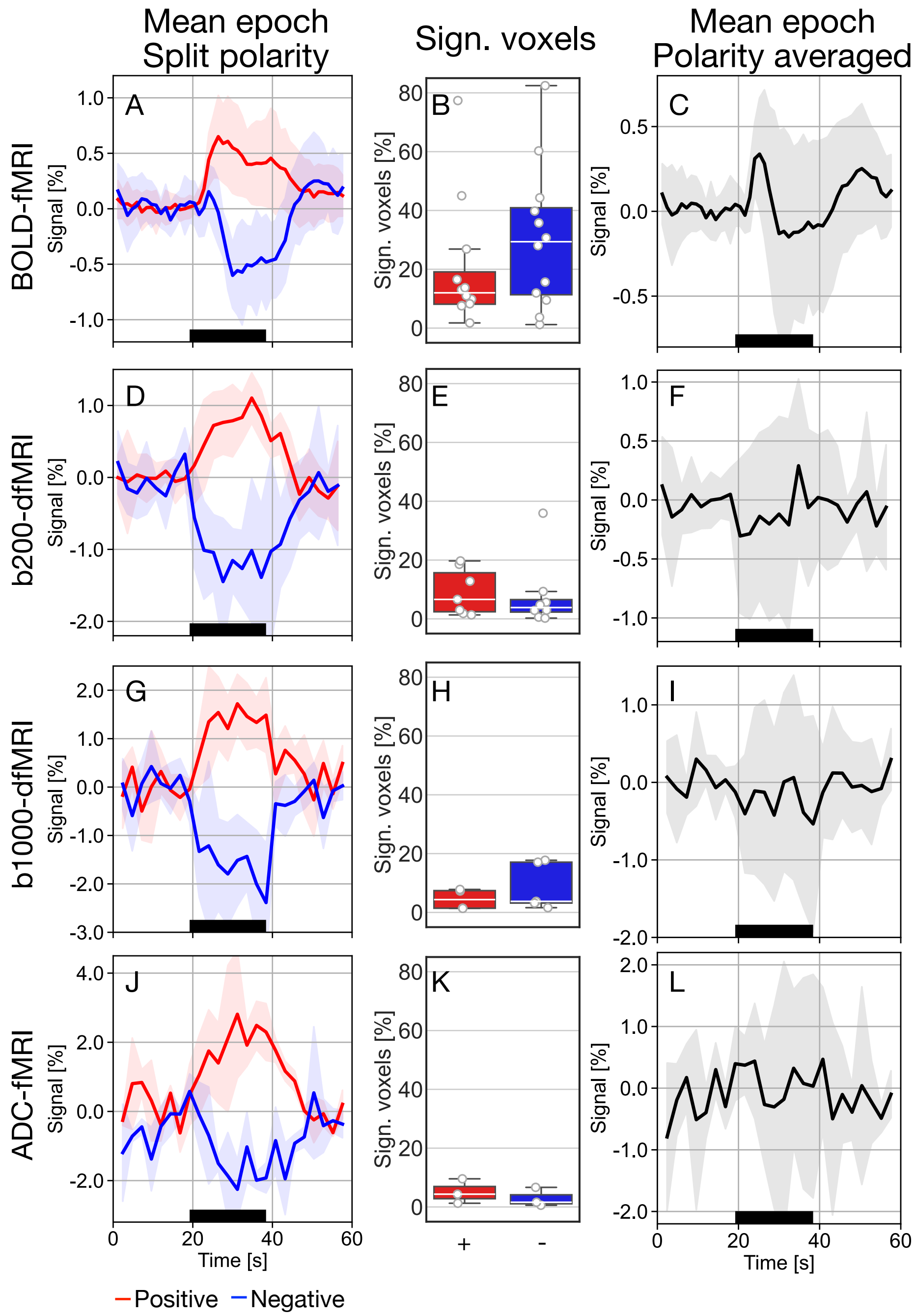


Supplementary Figure 10: **Mean response in significant voxels within SMA (HRF model).** Rows correspond to the BOLD-fMRI (n=12), b200-dfMRI, b1000-dfMRI, and ADC-fMRI (n=11) contrasts. Columns show mean epoch time courses in significant voxels within the supplementary motor area (SMA) split by polarity (left), corresponding percentage of significant voxels (middle), and mean epoch time courses averaged across polarities (right). Timeseries were first averaged across significant SMA voxels within each subject and then across subjects. Shaded grey areas indicate the standard deviation across subjects. TMS duration is indicated by black bars. Positive (+) and negative (-) associations are depicted in red and blue, respectively. Number of subjects contributing to the averaging: $n_{positive}$ = 12, 6, 4, 3, and $n_{negative}$ = 12, 6, 4, 3, for BOLD-fMRI, b200-dfMRI, b1000-dfMRI and ADC-fMRI, respectively. BOLD = blood oxygen level-dependent, ADC = apparent diffusion coefficient.
